# Bacteriophages targeting *Acinetobacter baumannii* capsule induce antimicrobial resensitization

**DOI:** 10.1101/2020.02.25.965590

**Authors:** Fernando Gordillo Altamirano, John H. Forsyth, Ruzeen Patwa, Xenia Kostoulias, Michael Trim, Dinesh Subedi, Stuart Archer, Faye C. Morris, Cody Oliveira, Luisa Kielty, Denis Korneev, Moira K. O’Bryan, Trevor J. Lithgow, Anton Y. Peleg, Jeremy J. Barr

## Abstract

Carbapenem-resistant *Acinetobacter baumannii* is responsible for frequent, hard-to-treat and often fatal healthcare-associated infections. Phage therapy, the use of viruses that infect and kill bacteria, is an approach gaining significant clinical interest to combat antibiotic-resistant infections. However, a major limitation is that bacteria can develop resistance against phages. Here, we isolated phages with activity against a panel of *A. baumannii* strains and focused on clinical isolates AB900 and A9844 and their phages for detailed characterization. As expected, coincubation of the phages with their hosts *in vitro* resulted in the emergence of phage-resistant bacterial mutants. Genome sequence analysis revealed that phage-resistant mutants harbored loss-of-function mutations in genes from the K locus, responsible for the biosynthesis of the bacterial capsule. Using molecular biology techniques, phage adsorption assays, and quantitative evaluation of capsule production, we established that the bacterial capsule serves as the primary receptor for these phages. As a collateral phenotype of impaired capsule production, the phage-resistant strains could not form biofilms, became fully sensitized to the human complement system, showed increased susceptibility to beta-lactam antibiotics, and became vulnerable to additional phages. Finally, in a murine model of bacteremia, the phage-resistant *A. baumannii* demonstrated a diminished capacity to colonize blood and solid tissues. This study demonstrates that phages can be used not only for their lytic activity but, if combined with *a posteriori* knowledge of their receptors and the mechanism of bacterial resistance, for their potential synergy with other antimicrobial agents, thus providing even broader clinical options for phage therapy.

## Introduction

In 2019, antimicrobial resistance was listed by the World Health Organization (WHO) as one of the top ten threats to global health (1). Multidrug resistant (MDR) infections are consistently associated with poor clinical outcomes and represent a significant financial burden on the healthcare system (2, 3). Also in 2019, the WHO and Centers for Disease Control and Prevention (CDC) prioritized carbapenem-resistant *Acinetobacter baumannii* as a pathogen critically requiring research and development of new antimicrobial strategies (4, 5). *A. baumannii*, a gram-negative coccobacillus, is a member of the *ESKAPE* group of pathogens, which are prominent for causing frequent and hard-to-treat healthcare-associated infections (6).

As a species, *A. baumannii* is highly resilient, and capable of surviving for months in biofilms on abiotic surfaces (7, 8). It can cause pneumonia, bacteremia, urinary tract infections, meningitis and wound infections, particularly in the context of intensive care units, and is frequently associated with indwelling medical devices (8). Previous studies have estimated *A. baumannii* to be the causative agent of up to 20% of infections within intensive care units worldwide (9). The pathogenicity of *A. baumannii* is facilitated by features of the bacterial cell surface, including secretion of an extracellular polysaccharide capsule, outer membrane proteome, and iron acquisition systems (10, 11). The extracellular capsule is of particular importance, given that it is an essential feature for biofilm formation (12), which in turn protects *A. baumannii* from the immune system, antibiotics and disinfectants (13, 14), and increases the risk of co-colonization with other pathogens (15). The capsule also allows the pathogen to escape the action of antibodies and complement-mediated killing, promotes intrinsic antimicrobial-resistant phenotypes, and increases virulence (16).

Bacteriophages (phages) are viruses that infect and kill bacteria, making them an attractive option to combat antimicrobial resistance (17). Phage therapy is the administration of phages directly to a patient with the purpose of lysing the underlying bacterial pathogen (18). Since phages are highly species-specific, phage therapy has a favorable safety profile and does not disturb the microbiome in the way antibiotics do. However, this also means that both diagnosis of the infection and identification of the etiological agent are required to generate a precise phage-host match that will maximize treatment effectiveness (17). Under these circumstances, it is useful to establish and expand collections of well-characterized phages against problematic pathogens such as *A. baumannii*. Phage therapy has shown promising results against *A. baumannii* in murine models of pneumonia (19), bacteremia (20) and wound infections (21). And in 2017, the first case of successful intravenous and intracavitary phage therapy targeting a systemic MDR infection was reported in the United States (22), involving a 68-year-old diabetic patient with necrotizing pancreatitis complicated by an MDR *A. baumannii* infection that was treated with successive administration of phage cocktails (22).

Phage specificity is regarded as a major limitation to making phage therapy more accessible. The primary determinant for phage specificity occurs during the first step of the phage infection cycle, which is the adsorption of the phage to its receptor on the bacterial cell surface. In the case of *A. baumannii*, the cell surface comprises a range of structures composed both of proteins and polysaccharides (11). The first layer of this cell surface are the capsular polysaccharides and, in *A. baumannii*, more than forty distinct capsular polysaccharide structures have been chemically determined and more than a hundred diverse capsule types predicted based on genome sequence analysis (16, 23). Likewise, the outer membrane proteome in *A. baumannii* can vary greatly from strain to strain (11). Given that there is a current lack of serotyping to provide information on the capsule-type of *A. baumannii* clinical isolates, experimentally determining a precise host-phage match is an important issue.

Bacteria rapidly adapt to external pressures and can evolve resistance to antibiotics and phages alike. Lytic phages impart strong selective pressures on their hosts, and phage-resistant phenotypes can emerge in clinically-relevant timeframes (24). In particular, *in vitro* phage resistance often arises through the modification or loss of surface receptors used during phage adsorption (25). This can result in the loss of the physiological function of the receptor, with a collateral reduction in bacterial fitness, often referred to as a ‘trade-off’ (26, 27). Trade-offs have been documented in various species and include decreased growth and impaired production of capsules in *Staphylococcus aureus* (28), loss of virulence in *Salmonella enterica* (29), and resensitization to antibiotics in *Pseudomonas aeruginosa* (30). Understanding phage-resistance mechanisms and their associated trade-offs can allow for their clinical exploitation, opening the door to the next generation of phage therapy.

In this study, we isolated and characterized novel phages active against clinical isolates of *A. baumannii*, including carbapenem-resistant isolates, to establish a phage library against this species. Upon examination of lytic phage activity, we observed that phage-resistant mutants quickly emerged *in vitro*. Using next-generation sequencing and bacterial genomics, we identified mutations in the genes associated with capsule production and confirmed that they were causative for the phage-resistant phenotypes. We used genetic engineering and phage adsorption assays to identify the capsule of *A. baumannii* as the primary phage receptor, and established that phage-resistance emerged through loss of this receptor and subsequent lack of phage adsorption. As a result, phage-resistant mutants exhibited altered capsule production, increased sensitivity to bactericidal agents, including human complement, antibiotics, and other phages, and reduced fitness in an *in vivo* bacteremia model. Here, we propose the use of phages, with *a posteriori* knowledge of their receptors, to enable both effective antimicrobial treatment and the informed prediction of phage-resistance outcomes, with exploitable fitness trade-offs extending the clinical impact of phage therapy.

## Results

### Characterization of *Acinetobacter baumannii-*specific phages

A group of nine *A. baumannii* strains, from different sites of infection and exhibiting different antimicrobial resistance profiles were selected for this study (31-35) (Fig. 1A). We used the Phage-on-Tap protocol (36) to isolate and purify phages into high-titer preparations targeting the nine strains. Eight bacteriophages were isolated and used to establish an *A. baumannii*-specific phage library. We tested the activity of the phages against all *A. baumannii* strains to determine the host infectivity range (Fig. 1A). The phage with the broadest host range, øCO01, was capable of productive infection in 4/10 strains, while phages øFG01, øCO02, øCO03 and øLK01 were highly specific and capable of productive infection in only their host of isolation.

**Figure 1.**
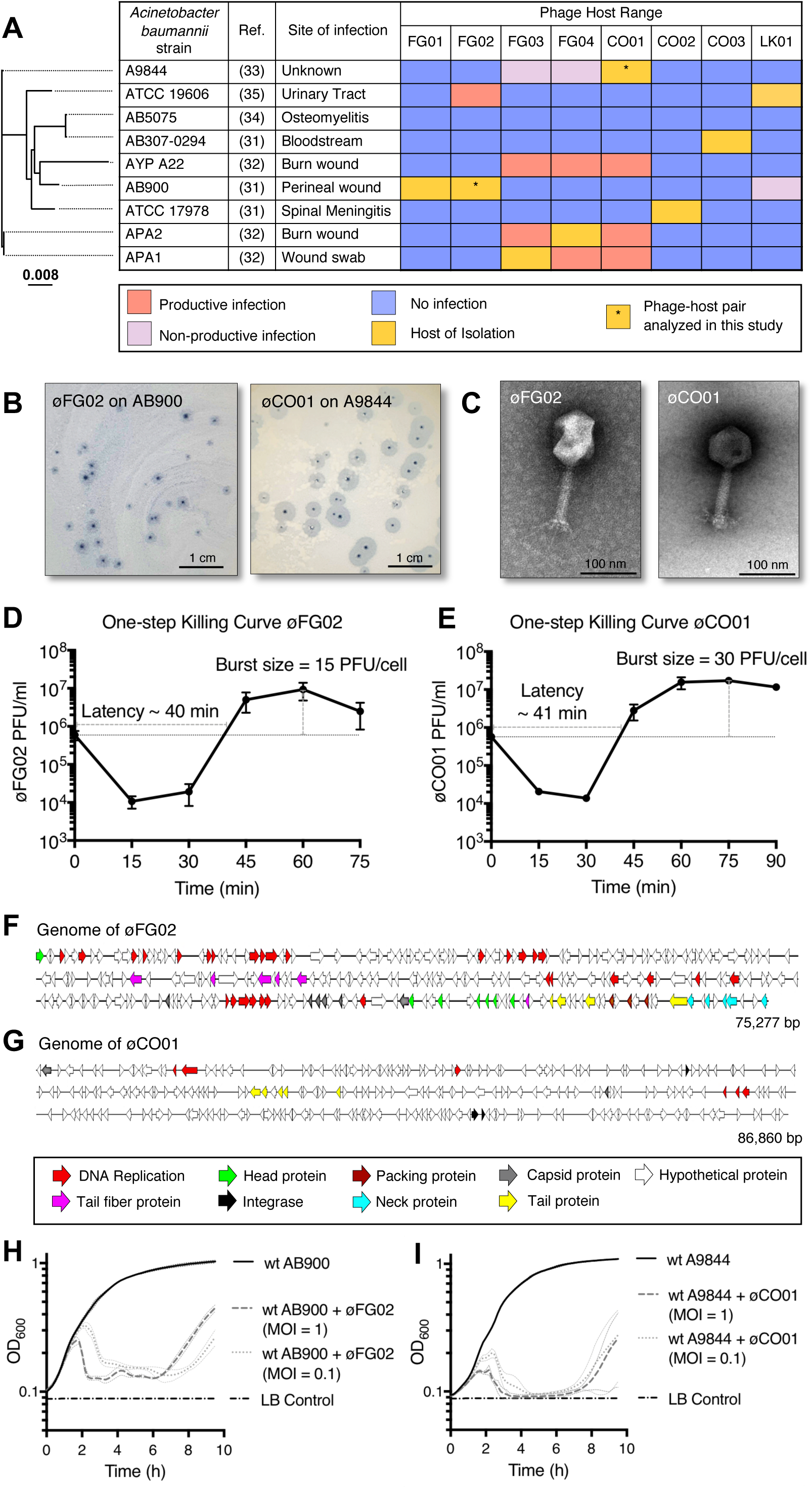
Characterization of *Acinetobacter baumannii-*specific phages. A: Host range of the eight *A. baumannii-*specific phages isolated in this study. Bacterial strains are ordered by the core-genome alignment phylogenetic tree on the left. Productive infection is defined as lysis on a spot assay and production of plaques on a soft agar overlay, whereas non-productive infection is defined as lysis on a spot assay with no plaques on a soft overlay agar. B: Plaque morphology of phage øFG02 on host AB900 and øCO01 on host A9844, showing the central clearings or lysis zones surrounded by hazy halos. C: Phage morphology as seen by transmission electron microscopy, øFG02 and øCO01 have icosahedral capsids and contractile sheathed tails, characteristic of the *Myoviridae* family. D and E: One-step killing curves of phages øFG02 and øCO01 on their hosts of isolation. Latency time was measured from the beginning of the experiment to when the curve at exponential growth reached the initial phage inoculum, whereas burst size was calculated by dividing the maximum free phage count after bacterial burst by the initial phage titer in the experiment. Error bars represent standard error of the mean (SEM) (n = 3). F and G: Genome assemblies of phages øFG02 and øCO01 show structural and replication genes for both. The genome of øCO01 shows the presence of three predicted integrases. H and I: Growth curves of *A. baumannii* strains AB900 and A9844 with and without phages. Phages were applied at two different multiplicities of infection (1 and 0.1). Lytic activity of phages is shown by the drop in optical density shortly after initiation of the bacterial exponential growth phase; and emergence of phage resistance is shown after 6 h of incubation. Each curve’s top and bottom dotted lines show the standard deviation (SD) (n = 3).

Phages øFG02 and øCO01 were chosen for further characterization based upon their contrasting host ranges, high titers achievable during experimental amplification, and large phylogenetic distance between their hosts of isolation (AB900 and the carbapenem-resistant isolate A9844, respectively). Plaque morphology of both phages presented a distinctive hazy halo surrounding a zone of clear lysis (Fig. 1B). Microscopic examination of the phages revealed virion lengths of ∼250 nm for øFG02 and ∼200 nm for øCO01, with icosahedral capsids and sheathed contractile tails, consistent with the *Myoviridae* family of bacteriophages (Fig. 1C). The one-step killing curve of øFG02 demonstrated a latency period of approximately 40 min, with a burst size of 15 plaque-forming units (PFU) per infection (Fig. 1D). For øCO01, the latency was ∼41 min but the burst size was doubled, at 30 PFU/infection (Fig. 1E).

Genome sequencing of øFG02 generated a single contig of 75,277 bp (44.4% G+C), which best matched with *Cronobacter* phage vB_CsaM_leN (90% coverage, 91.62% identity, GenBank accession number KX431560.1). RAST annotation predicted 255 protein-coding regions, and the VIBRANT (37) analysis—which uses multiple databases to annotate, determine the completeness and characterize virome function—revealed that øFG02’s genome possessed a lytic nature, and predicted several phage structural and replication genes (Fig. 1F). For phage øCO01, we identified a contig of 86,860 bp (37.5% G+C) with a best match against *Acinetobacter* phage Ab105-3phi (54% coverage and > 89% identity, GenBank accession number KT588073.1). However, the contig also had high similarity (> 99% coverage and > 80% identity) with several *A. baumannii* genomes from the NCBI database and the VIBRANT analysis was negative for a lytic feature. Annotation of the contig revealed integrase genes along with phage structural genes (Fig. 1G). Based on this finding, we attempted to experimentally produce A9844 lysogens containing øCO01 via coincubation. We performed the experiment with two different concentrations of øCO01, and screened 18 colonies from each, without success (Fig. S1). This suggests that despite the presence of a predicted integrase in the genome of øCO01, it is incapable of lysogenizing its host of isolation, at least under regular *in vitro* conditions.

To further demonstrate the lytic activity of the phages, we coincubated them with their bacterial hosts at two different multiplicities of infection (MOI), 1 and 0.1. The growth curves (Fig. 1H and 1I) show phage lytic activity beginning during the bacterial exponential growth phase, with the bactericidal effect occurring earlier at a higher MOI. Both phages inhibited bacterial growth for just over 6 h and shortly thereafter, regardless of the phage MOI used, the strains developed phage-resistance and resumed growth *in vitro*.

### Isolation of phage-resistant *A. baumannii* mutants

Bacteria are known to quickly develop phage-resistance when subjected to the selective pressure of lytic phages *in vitro*. We coincubated strains AB900 and A9844 in a semi-solid culture with phages øFG02 and øCO01, respectively. Colonies that were able to grow in the center of lysis zones (Fig. S2) subsequently underwent two single-colony isolation steps. Next, we performed inverted spotting assays (Fig. 2A and 2B) and growth curves (Fig. 2C and 2D) to confirm the phage-resistant phenotype of the selected colonies. From these assays, we picked a single mutant of strain AB900 that was resistant (R) to phage øFG02 (øFG02-R AB900) and a single mutant of strain A9844 that was resistant (R) to phage øCO01 (øCO01-R A9844). *In vitro*, phage-resistant mutants did not revert to the wild type phenotype despite ≥ 10 growth cycles (defined as the subculture on a liquid or solid medium without the presence of phage). Growth curves showed a 3% increase in area under the curve (AUC) for øFG02-R AB900 and a 9% decrease for øCO01-R A9844 when compared to their wild type counterparts (Fig. S3).

**Figure 2.**
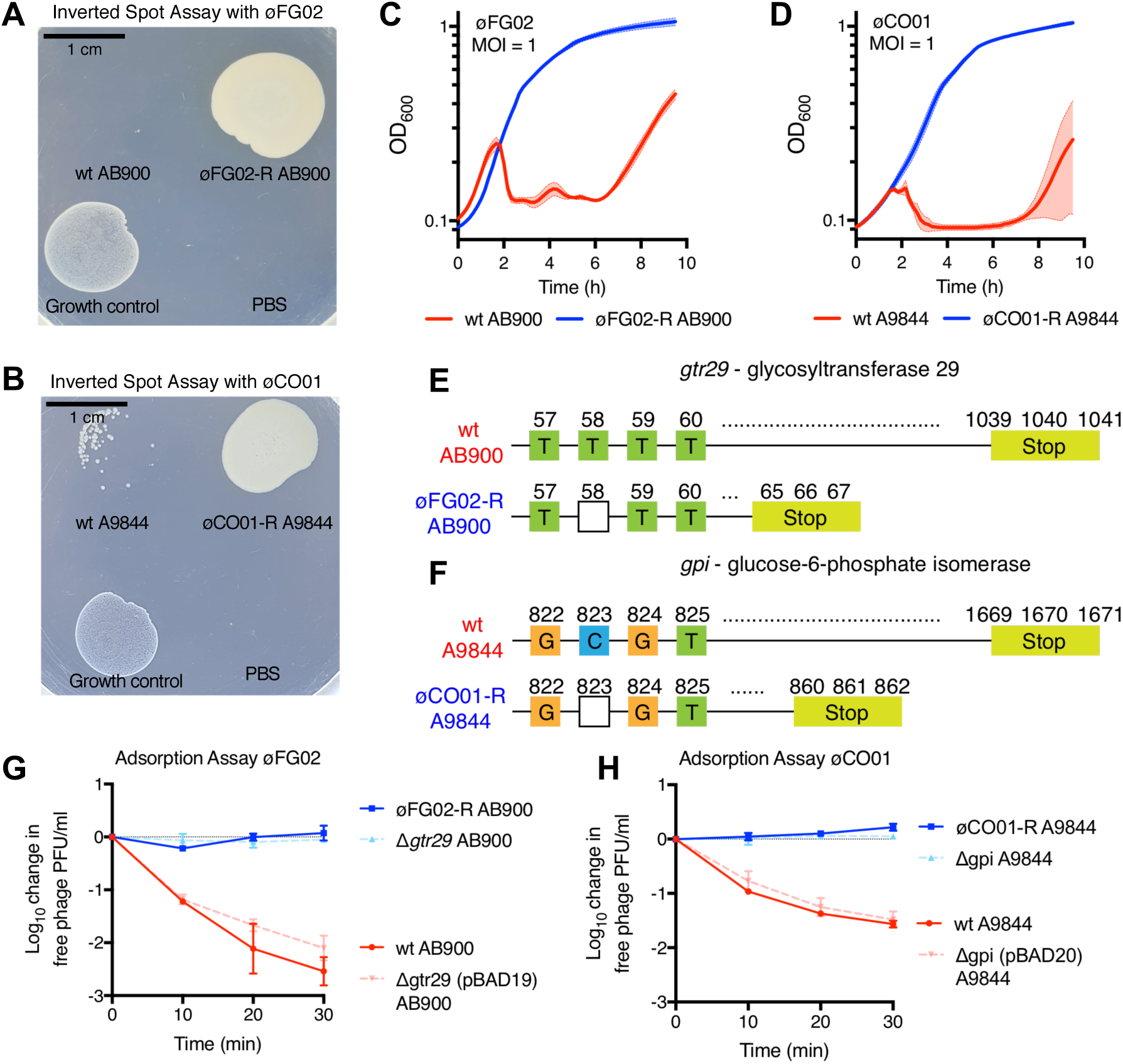
*A. baumannii* phage-resistant mutant strains, phage receptors and mechanism of phage resistance. A and B: Inverted spot plate assay containing 10^8^ PFU of phages øFG02 or øCO01. Lack of confluent growth of AB900 and A9844, and confluent growth of øFG02-R AB900 and øCO01-R A9844 suggest phage-resistant phenotype. PBS used as a negative control, and *Enterococcus faecium* as a bacterial growth control. C and D: Growth curves in the presence of phages at an MOI of 1. Wild type strains AB900 and A9844 (in red) were affected by phage activity, whereas phage resistant mutants øFG02-R AB900 and øCO01-R A9844 (in blue) grew unaffected. Shaded zones represent SD (n = 3). E and F: Mutations identified in øFG02-R AB900 and øCO01-R A9844 associated with the phage-resistant phenotype. For both strains, the affected genes are in the K locus, and the single-nucleotide deletions led to premature truncation of the proteins. G and H: Phage adsorption assay, log_10_ reduction in free phage titers over a 30 min interval after mixing phages and hosts. Each phage was mixed with either wild type, phage-resistant, knockout mutant, or plasmid-complemented hosts. Phages adsorb to phage-sensitive strains (wild type and plasmid-complemented, shades of red) leading to a decrease in free phage titer, whereas phages do not adsorb to phage-resistant strains (phage-resistant, knockout mutant, shades of blue). Error bars represent SEM (n = 3).

### Identification of phage receptors and phage-resistance mechanisms in *A. baumannii*

With the purpose of identifying the number, type, and loci of the mutations granting phage-resistance to our *A. baumannii* strains, we extracted genomic DNA from the wild type strains AB900, A9844 and their phage-resistant counterparts øFG02-R AB900 and øCO01-R A9844. Illumina® HiSeq 150 bp paired-end reads were used for a pairwise comparison of the bacterial genomes. In each phage-resistant strain, a single nucleotide deletion in a gene within the K locus was found (Fig. 2E and 2F). Genes in the K locus of *A. baumannii* regulate the production, modification and export of capsular polysaccharides (38). For øFG02-R AB900, the affected gene was *gtr29*, which codes for a glycosyltransferase and in øCO01-R A9844 it was *gpi*, which codes for the enzyme glucose-6-phosphate isomerase. In both cases, the identified deletions caused codon frameshifts resulting in the premature truncation of the proteins, at amino acid 21 out of 346, and 286 out of 556 for the glycosyltransferase and glucose-6-phosphate isomerase, respectively (Fig. 2E and 2F).

To demonstrate that the identified mutations were responsible for the phage-resistant phenotypes, we disrupted genes *gtr29* and *gpi* in AB900 and A9844, respectively, using a kanamycin-resistance cassette. Next, we used the newly-obtained strains Δgtr29 AB900 and Δgpi A9844 in an efficiency of plating (EOP) assay and confirmed they were completely resistant to phages øFG02 and øCO01, respectively (EOP = 0%). Complementation analysis showed restoration of phage susceptibility (EOP = 100%), thus confirming the involvement of genes *gtr29* and *gpi* in phage infectivity.

Both *gtr29* and *gpi* are in the K locus and thus predicted to function in biosynthesis of capsular polysaccharides. We hypothesized that phages øFG02 and øCO01 used capsular polysaccharides as their receptors, and that the loss or modification of such receptors led to lack of phage adsorption and subsequent phage-resistance in *A. baumannii*. To test the hypothesis, we performed an adsorption assay using each phage and their respective strain set (wild type, phage-resistant isolate, knockout mutant and plasmid-complemented mutant) (Fig. 2G and 2H). Within 30 min, more than 99% and 97% of øFG02 and øCO01 particles had adsorbed to their respective wild type hosts. In contrast, the titers of free phage particles did not decrease when the phages were mixed with either the phage-resistant or knockout mutants, demonstrating the inability of phages to adsorb to these bacterial cells. As expected, phage adsorption was reinstated in the plasmid-complemented strains expressing *gtr29* and *gpi.* The adsorption assay supported our hypothesis that capsular polysaccharides are the phage receptor for øFG02 and øCO01 and that phage-resistance arose through alteration or loss of these receptors leading to inhibition of phage adsorption.

### *A. baumannii* phage-resistant strains exhibit phenotypic trade-offs

As a next step, we sought to investigate if our phage-resistant *A. baumannii* strains would exhibit detrimental phenotypic trade-offs associated with their acquired mechanism of phage-resistance. We began tackling this question by assessing the production of capsular polysaccharides in the phage-resistant strains.

First, we used Maneval’s staining technique to visually evaluate capsule production (Fig. 3A). Capsules were observed as unstained halos surrounding each bacterial cell (stained in pink), and the thickness of such halos is a proxy of capsule production. Wild type strains AB900 and A9844 were uniformly capsulated, whereas most of the cells of øFG02-R AB900 and øCO01-R A9844 appeared to lack capsules. Single cell imaging using scanning electron microscopy (SEM) (∼100 cells per strain observed, consistent phenotypes throughout the observations), suggested that the surface of wild type A9844 possessed a complex polysaccharide capsule, including many pilus-shaped protrusions. In contrast, the appearance of the phage-resistant surface suggested loss of capsule. Less obvious differences were noted between AB900 and øFG02-R AB900 (Fig. 3B and S4). For a quantitative assessment of the phenotype, we assayed the production of surface polysaccharides using the colorimetric sulfuric acid and phenol assay (39) (Fig. 3C). Using this technique, we observed 2.2-fold and 3-fold decrease in polysaccharide production in øFG02-R AB900 and øCO01-R A9844, respectively, compared to their wild type strains (n = 3; 79 ± 1.7 μg/ml vs. 35.6 ± 8.8 μg/ml [mean ± SD]; unpaired t test; p = 0.001; two-tailed; for AB900 vs. øFG02-R AB900) (n = 3; 179.9 ± 15.6 μg/ml vs. 58.1 ± 5.6 μg/ml; unpaired t test; p = 0.0002; two-tailed; for A9844 vs. øCO01-R A9844). In accordance, colonies of strains AB900 and A9844 exhibited a mucoid phenotype—viscous and sticky colonies—characteristic of capsule production, which was lost in colonies of øCO01-R A9844 but not in øFG02-R AB900. Taken together, our findings indicate a major defect in the overall production of capsule polysaccharides in the phage-resistant mutant strains, with these results being consistently more pronounced in øCO01-R A9844.

**Figure 3.**
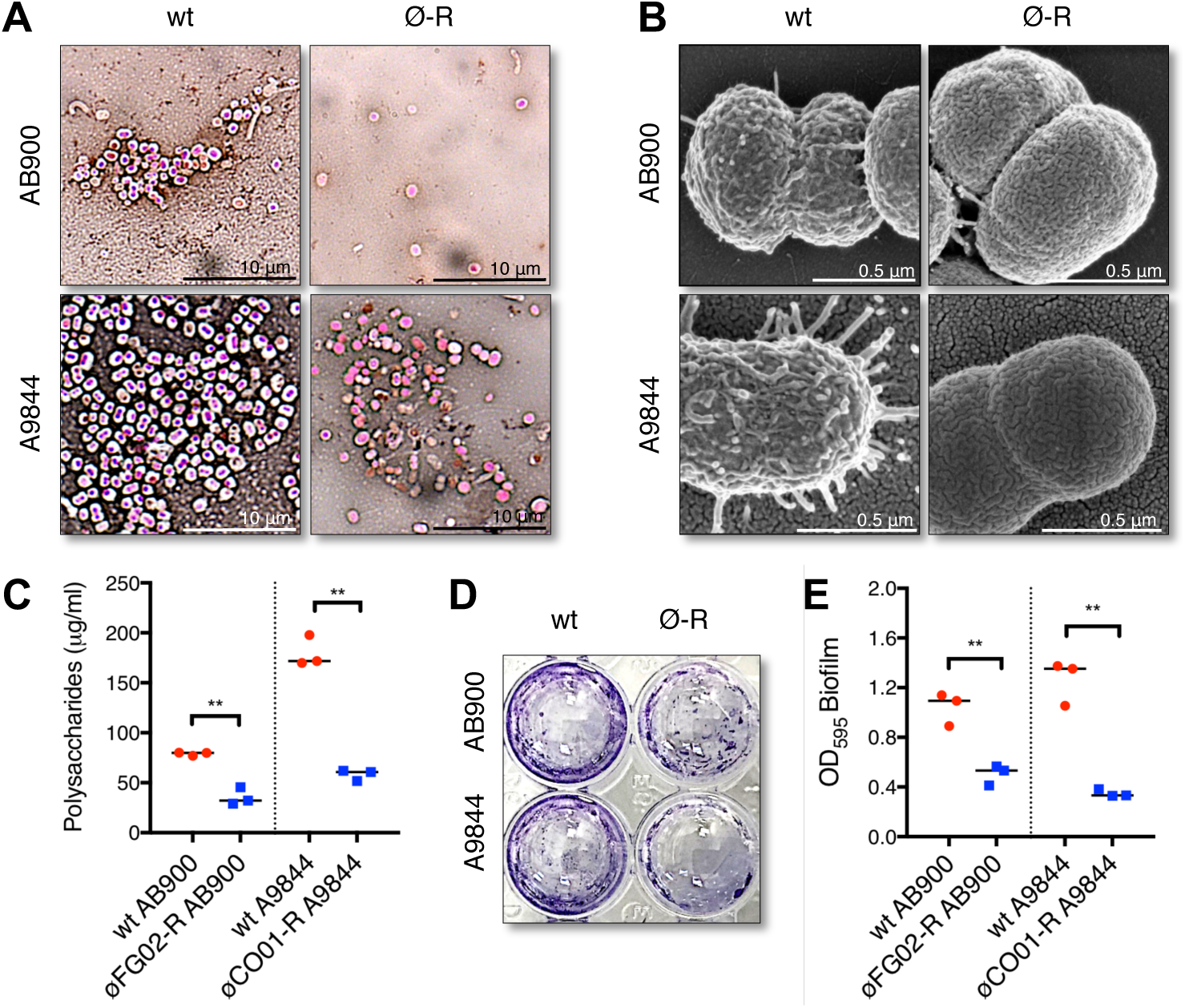
Phenotypic trade-offs of phage-resistance in *A. baumannii.* A: Microscopic inspection of capsule thickness in slides stained using the Maneval’s technique. B: Cell surface appearance as observed via scanning electron microscopy, representative images from ∼100 observations (additional images in Fig. S4). C: Production of capsule polysaccharides measured by absorbance of a colorimetric phenol-sulfuric acid reaction and extrapolated to a standard curve (Fig. S5). D and E: Biofilm production on a polystyrene surface at 48 h, measured by absorbance of crystal-violet stained and ethanol-solubilized biofilm. For the scatterplots in panels C and E, wild type strains represented in red and phage-resistant mutants in blue, bars represent medians (n = 3), each point represents the average from three technical replicates, unpaired t test ** = p < 0.005, two-tailed.

Capsular polysaccharides aid in the attachment of bacterial cells to biotic and abiotic surfaces, which is the first step in the formation of biofilms (16), one of the hallmarks of *A. baumannii* clinical infections. Having observed a significant decrease in capsule production in the phage-resistant mutants, we tested if these mutants had impaired biofilm production. We compared the ability of the strains to form biofilms on a polystyrene surface at 48 h (Fig. 3D). As expected, the phage-resistant strains øFG02-R AB900 and øCO01-R A9844 had a 2.1-fold and 3.6-fold reduction, respectively, in biofilm production when compared to their wild type counterparts (Fig. 3E) (n = 3; optical density of biofilm 1 ± 0.1 vs. 0.5 ± 0.1 [mean ± SD]; unpaired t test; p = 0.0038, two-tailed; for AB900 vs. øFG02-R AB900) (n = 3; 1.3 ± 0.2 vs. 0.4 ± 0.03; unpaired t test; p = 0.001, two-tailed; for A9844 vs. øCO01-R A9844). Thus, the acquisition of phage-resistance in our *A. baumannii* strains resulted in the phenotypic trade-offs of impaired capsule production and biofilm formation.

### Phage-resistant *A. baumannii* strains are more susceptible to antimicrobial agents

Capsules and biofilms are major virulence determinants in *A. baumannii.* Part of their importance in bacterial pathogenesis relies on enabling *A. baumannii* to evade components of the host’s immune response and to resist the effect of several antibiotics. Following the observed trade-offs of our *A. baumannii* phage-resistant mutants, we hypothesized the mutants would have become vulnerable to the action of various antimicrobial agents.

The complement system consists of a cascade of serum proteins capable of eliminating pathogens by attacking their membrane, and enhancing the activity of antibodies and phagocytes. We established a serum killing assay to determine the susceptibility of *A. baumannii* to the action of the human complement system (Fig. 4A). Over a period of 180 min, wild type strains AB900 and A9844 withstood the effect of, and even grew in, human serum. Conversely, the phage-resistant mutants øFG02-R AB900 and øCO01-R A9844 were reduced below the level of detection in the serum even at the first timepoint of 45 min, representing a >4-log decrease. The results shown in the experiment can be safely attributed to the action of the complement system, as the antimicrobial effect was lost if heat-inactivated serum was instead used (data not shown).

**Figure 4.**
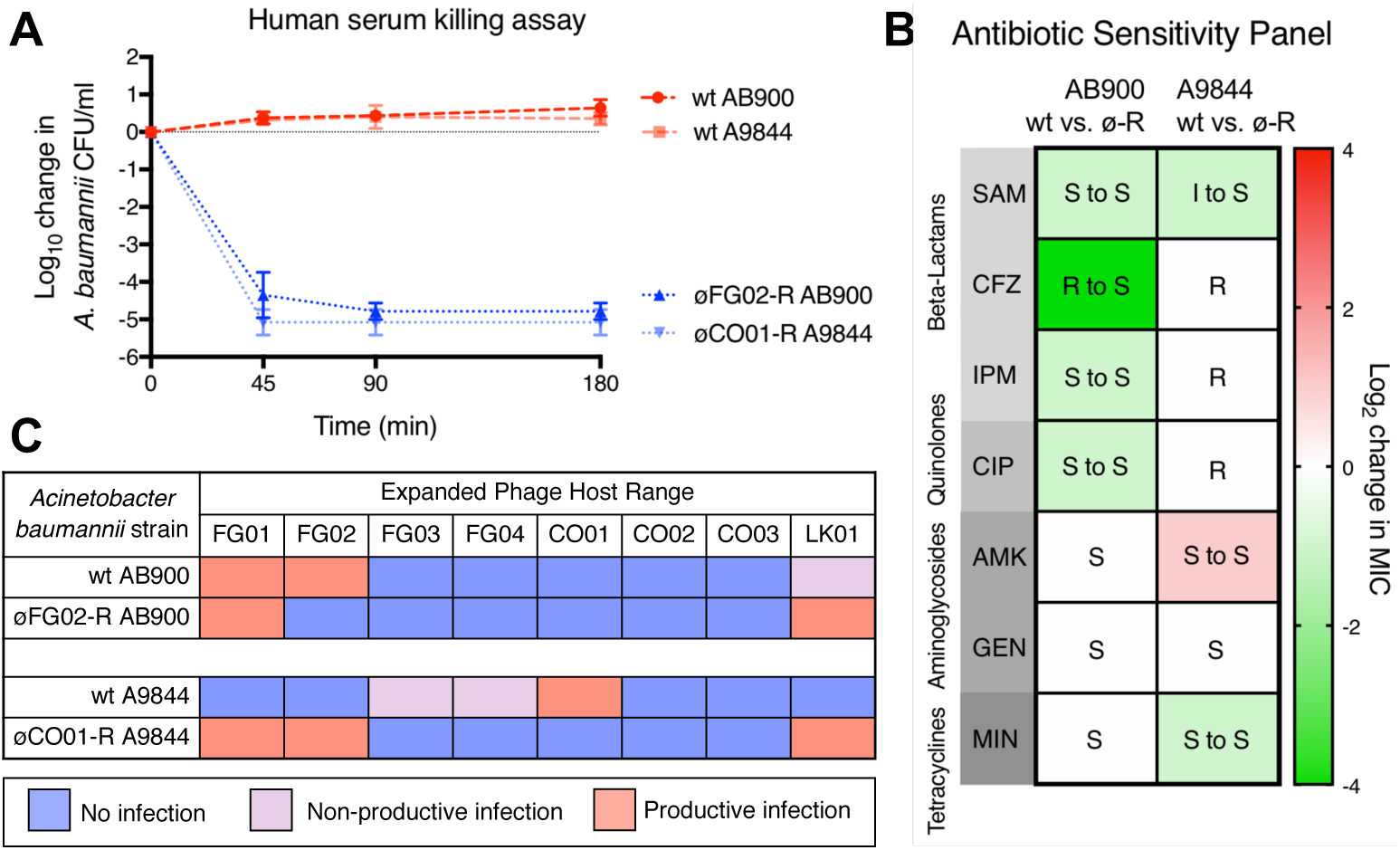
Increased sensitivity of phage-resistant *A. baumannii* to antimicrobial agents. A: Human serum killing assay. Freshly-thawed human serum was inoculated with 10^5^ CFU/ml of *A. baumannii*, and the change in bacterial load was measured at regular intervals. Wild type strains (shades of red) grew in serum whereas phage-resistant strains (shades of blue) were rapidly killed; error bars represent SEM (n = 3). B: Antibiotic sensitivity pattern. The minimum inhibitory concentrations (MICs) of 7 antibiotics, from 4 pharmacological groups were measured using the microbroth dilution method. The median (n = 3) of the log_2_ change in MIC between the wild type and phage-resistant strains is color coded, with shades of green showing a reduction in MIC in phage-resistant mutants, shades of red showing an increase in MIC, and white showing no change for each specific antibiotic. Inside each cell, the clinical interpretation of the MIC is shown (S = Sensitive, I = Intermediate, R = Resistant). Raw data values are available in Table S1. SAM: ampicillin + sulbactam, CFZ: ceftazidime, IPM: imipenem, CIP: ciprofloxacin, AMK: amikacin, GEN: gentamicin, MIN: minocycline. C: Expanded phage host range map of the *A. baumannii*-specific phage library against øFG02-R AB900 and øCO01-R A9844 reveals sensitization to additional phages.

As our mutants demonstrated heightened sensitivity to complement, we next sought to determine whether their susceptibility to seven clinically relevant antibiotics was also impacted. We used the microbroth dilution method and the CLSI guidelines for clinical interpretation (Fig. 4B, raw data in Table S1). Although AB900 is not an MDR strain, it is resistant to ceftazidime, a third-generation cephalosporin. The phage-resistant mutant strain øFG02-R AB900 presented a 16x decrease in the minimum inhibitory concentration (MIC) of ceftazidime, becoming sensitive to it, as well as 2x decrease in the MIC of fellow beta-lactams ampicillin + sulbactam and imipenem, and the fluoroquinolone ciprofloxacin. In contrast, A9844 is an MDR, carbapenem-resistant isolate, considered by the CDC as a priority pathogen. We observed that øCO01-R A9844 presented a 2x reduction in the MIC of minocycline and ampicillin + sulbactam. The latter resulted in a change of the clinical interpretation of the MIC, making A9844 sensitive to ampicillin + sulbactam. However, øCO01-R A9844 also exhibited a 2x increase in the MIC for the aminoglycoside amikacin, but without a change in the clinical interpretation. The findings show instances of antibiotic resensitization occurring as a trade-off of phage-resistance in *A. baumannii.*

Finally, we tested another antimicrobial strategy, our *A. baumannii-*specific phage library, against strains øFG02-R AB900 and øCO01-R A9844 for extended phage-host range (Fig. 4C). We observed that, upon acquiring resistance to phage øFG02, strain øFG02-R AB900 became sensitive to phage øLK01, which was previously only capable of non-productive infection in wild type AB900. Furthermore, strain øCO01-R A9844 became vulnerable to the lytic action of three new phages, øFG01, øFG02 and øLK01. In summary, acquisition of phage-resistance was accompanied by resensitization to at least three different types of antimicrobial agents. This discovery is relevant, as all of the agents can play a role in the clinical setting during the treatment of *A. baumannii* infections.

### Phage-resistant *A. baumannii* strains show decreased fitness *in vivo*

The trade-offs of impaired capsule production and resensitization to antimicrobials could affect *A. baumannii*’s ability to invade and survive in mammalian hosts (40, 41). We therefore sought to observe the *in vivo* fitness of our phage-resistant mutants. To achieve this, we established a murine model of septicemia via intraperitoneal injection of 10^6^ CFU with 6% porcine stomach mucin. Groups of 6-to-10 weeks old, female, BALB/c mice (n = 4) were infected with either AB900, A9844, or their phage-resistant counterparts. After 8 h, the levels of bacterial colonization were assessed for the blood, liver, kidney and spleen. In both strain pairs, we observed a 2-log reduction of bacterial load in solid organs and a >3-log reduction in blood for phage-resistant mutants, when compared to wild type strains (Fig. 5A and 5B) (n = 4; Mann-Whitney test, p = 0.0286 for all comparisons except kidney in A9844 [p = 0.0571], two-tailed). Interestingly, after their passage through the animal hosts, øCO01-R A9844 bacteria did not revert to the wild type form, but in 2 out of the 4 mice infected with øFG02-R AB900 there was evidence of reversion to wild type AB900. This reversion was only seen in bacteria retrieved from solid organs and not blood and, in one of the animal hosts, it was heterogeneous, meaning that populations of both wild type-reverted and phage-resistant mutants were retrieved. These observations supported the hypothesis that phage-resistant *A. baumannii* suffers a fitness cost *in vivo*.

**Figure 5.**
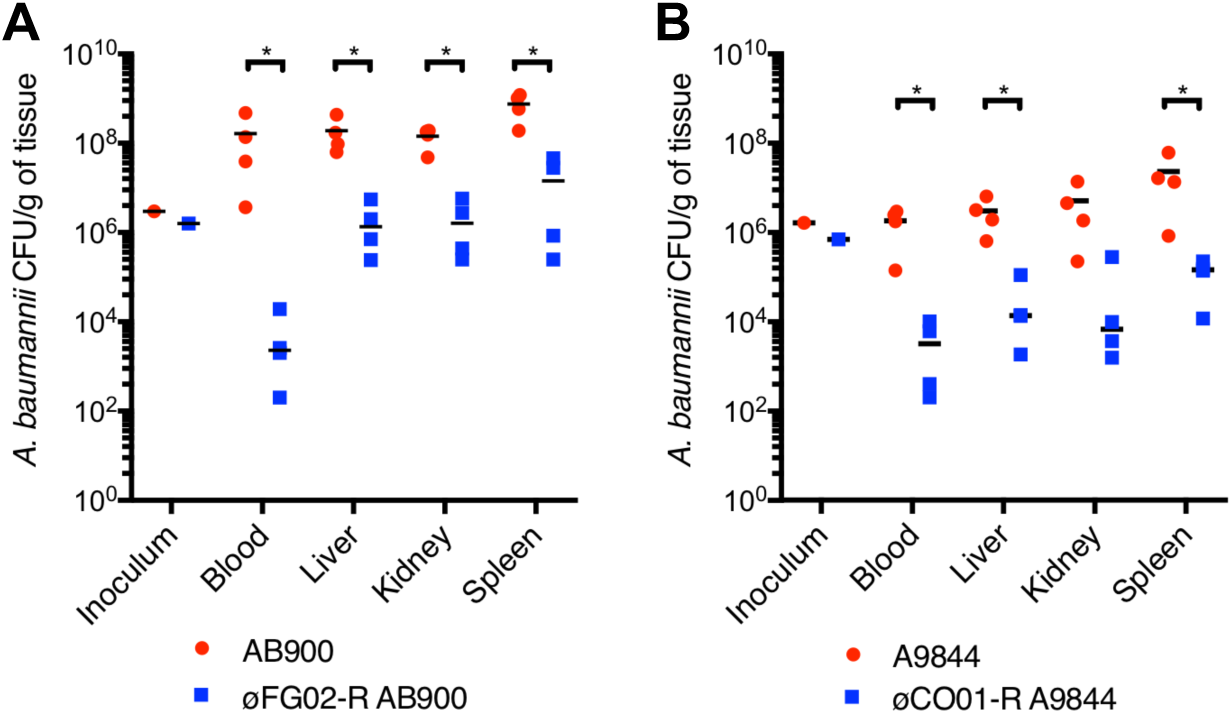
*In vivo* fitness is reduced in phage-resistant *A. baumannii.* 4 groups of 4 female, 6-to-10 weeks old, BALB/c mice were subjected to intraperitoneal (IP) injection of 10^6^ colony forming units of either AB900, A9844, øFG02-R AB900 or øCO01-R A9844. At the humane endpoint of 8 h, blood, liver, kidney and spleen were collected for bacterial quantification. A and B: Bacterial burdens at 8 h post infection, normalized by tissue weight. Wild type strains in red, phage-resistant strains in blue. Each data point represents an animal, with black bars representing the median values, Mann-Whitney test * = p < 0.05, two-tailed.

## Discussion

Antibiotic-resistant *A. baumannii* was recently deemed the top priority pathogen requiring new therapeutic strategies (4), and it certainly is one of the most threatening pathogens encountered in clinical practice (8). In this study, we isolated novel phages against a phylogenetically diverse panel of *A. baumannii* strains and provided a detailed characterization of two of them: øFG02 and øCO01. Despite the strong lytic activity of these phages, we observed the rapid emergence of phage-resistant mutants *in vitro*. Phage-resistance is often seen as a prominent road-block for efficacious phage therapy, with up to 80% of preclinical and clinical phage therapy trials reporting emergence of phage-resistant mutants (42). Detailed characterization of phage biology and identification of host receptors enabled us to identify the mechanism of phage-resistance in this pathogen. Phage-resistant strains harbored loss-of-function mutations in genes of the K locus, which are responsible for the biosynthesis of capsular polysaccharides. Using genetic engineering, we confirmed that disruption of glycosyltransferase 29 and glucose-6-phosphate isomerase led to phage-resistance in *A. baumannii* strains AB900 and A9844, respectively. Furthermore, an adsorption assay supported our hypothesis that phage-resistance emerged through disruption the phage receptors resulting in lack of phage adsorption. We also determined that capsule loss in phage-resistant *A. baumannii* is associated with a reduction in biofilm formation, resensitization to antimicrobial agents including antibiotics, human complement, and additional phages, and a reduction in fitness in a murine model of bacteremia. These results demonstrate that the capsular polysaccharides of *A. baumannii* are the receptors for phages øFG02 and øCO01. To the best of our knowledge, this is the first confirmation of a phage receptor in this key MDR superbug. These findings provide a step towards the exploitation of the fitness trade-offs of phage-resistance and improvement in the clinical translation of phage therapy.

The therapeutic use of lysogenic phages is subject to ongoing debate. Their use is normally inadvisable due to risks such as lysogenic conversion of the host, or reduced clinical effect (17). However, in the absence of lytic phages against a particular pathogen, or in emergency cases, their use might be justified (17). The genomic analysis of one of our phages, øCO01, revealed three putative integrase genes and significant sequence homology with *A. baumannii* genomes, suggesting a possible lysogenic nature. Although we cannot conclusively rule out the possibility of temperate behavior in øCO01, the phage displayed strong lytic activity in phenotypic assays (Figs. 1B, 1E and 1I), and was not able to lysogenize its host of isolation, A9844, *in vitro* (Fig. S1). Notably, the sequence analysis of øCO01 revealed a best match to *Acinetobacter* phage Ab105-3phi with a modest 54% coverage and >89% identity, so further investigation of gene function in øCO01 is warranted. It is possible that øCO01 emerged through recombination between a lytic and a lysogenic phage, and while it contains putative integrases, it behaves as a lytic phage on host A9844 under *in vitro* conditions. We suggest that experimental validation of a phage’s capacity to lysogenize a host, rather than simply the presence of a putative integrase gene, may be a more appropriate determinant for the selection of phages for therapeutic use.

Bacterial capsules have been demonstrated to be key virulence factors (11, 16). In *A. baumannii*, capsule biosynthesis is a complex and well-regulated process that involves up to 10 different groups of proteins (16, 38). The capsule is comprised of tightly packed repeating subunits (K units) that consist of 4 to 6 sugars, with a large diversity of K unit structures identified to date (16, 23). From our phage-resistant *A. baumannii* isolates, we identified mutations in two enzymes that are required for the early stages of K unit production. Specifically, the enzyme glucose-6-phosphate isomerase catalyzes the conversion of glucose-6-phosphate to fructose-6-phosphate, forming one of the many simple sugars that can constitute part of a K unit, while the glycosyltransferase is required to assemble specific sugars into the growing chain of the K unit. The biochemical structures of the K units of the capsules of AB900 and A9844 have been previously described (23, 43, 44). However, the exact effects that our mutations had on these structures were not elucidated here and were outside the scope of this study. As a result, we were only able to conclude that phages øFG02 and øCO01 use the capsule polysaccharides of *A. baumannii* as receptors, a phenomenon previously described for phages targeting the pathogens *Campylobacter jejuni* (45), *Salmonella enterica* (46) and *Klebsiella pneumoniae* (47), with the specific K unit polysaccharide structure remaining to be identified.

The most clinically important trade-off of phage-resistance observed in our *A. baumannii* strains was perhaps the resensitization to three types of antimicrobial agents. Antibiotic resensitization could be broadly explained through loss of capsule, theoretically making it easier for the drugs to access their target site. However, the fact that resensitization primarily occurred towards beta-lactams—which act via disruption of the cell wall—suggests a more specific mechanism. It has been described, for example, that inactivating mutations in genes encoding the Wzc or Wza components of the capsule secretion system in *A. baumannii* also lead to increased susceptibility to beta-lactams (11). It is possible that capsule-deficient bacterial cells have a lower tolerance threshold to further disruptions in components of the cell envelope. Further research is needed to determine if antibiotic protection provided by *A. baumannii* capsules is strain specific, capsule-type specific, or universal (16). It is also worth noting that despite the microbroth dilution method being the gold standard for antibiotic sensitivity testing (48), we cannot rule out the influence of test variability in our results, particularly in changes in MIC ≤ 2x between wild type and phage-resistant mutants. Susceptibility to complement-mediated killing can also be explained by the loss of capsule, as the hydrophilicity and negative charge of capsule sugars would normally limit complement deposition on the bacterial surface (11). Finally, the alteration of the surface polysaccharides in the phage-resistant mutants could have unmasked or generated other receptors that were recognized by additional phages, explaining the observed changes in the phage host range, and allowing us to enhance and extend the use of phage therapy. Interestingly, both phages analyzed in this study presented plaque morphologies of lytic centers surrounded by hazy halos (Fig. 1B), which suggest the production of depolymerases, enzymes capable of digesting capsular polysaccharides (49). Further research could pursue the purification and characterization of the biological activity of these enzymes, attempting to replicate the described effects of capsule loss in *A. baumannii.*

Despite being situated within the K locus in *A. baumannii*, the physiological role of glycosyltransferase 29 and glucose-6-phosphate isomerase may go beyond capsule production. Genes from the K locus are also used during the biosynthesis of outer membrane lipooligosaccharides (16), and some of the proteins they encode have been described as “multitaskers” involved in bacterial virulence and metabolism (50). As such, it is possible that further effects of the disruption of *gtr29* and *gpi*, unrelated to capsule loss, could be identified in our phage-resistant mutants. Altered production of glycosyltransferases and glucose-6-phosphate isomerase has been previously shown to affect virulence and resistance to antimicrobials through different routes in a few pathogens. Boinett et al. (51) identified *gpi* and at least two glycosyltransferases as some of many upregulated genes in colistin-resistant strains of *A. baumannii.* Zhang et al. (52) demonstrated that isomerase-deficient mutants of the fungal pathogen *Cryptococcus neoformans* had reduced capsule biosynthesis, disruptions in their cell wall and plasma membrane and exhibited hypersensitivity to osmotic stress. Moreover, Tsuge et al. (53) noted a significant reduction in virulence upon inactivation of glucose-6-phosphate isomerase in the plant pathogen *Xanthomonas oryzae*, possibly due to impaired carbohydrate uptake. Investigating further trade-offs in our phage-resistant mutants could lead to the discovery of other clinically exploitable phenotypes.

Our results suggest that the capsule of *A. baumannii* acts as a phage receptor and tropism determinant. We worked with two *A. baumannii* phages isolated from phylogenetically distant strains, both of which induced loss-of-function mutations: although distinct, the affected genes were located within the same locus and are both involved in the production of capsular polysaccharides. This is further supported by recent studies documenting that phage tail fibers directly interact with *A. baumannii* capsular polysaccharides (54, 55) and a phage therapy case study treating a complicated MDR *A. baumannii* infection that reported capsule loss upon emergence of phage-resistance (22). However, it should be noted that all these studies have focused on a small number of phages and that other *A. baumannii* phage receptors are likely.

We attribute the emergence of phage-resistance to the loss of *A. baumannii*’s capsule, which subsequently results in clinically exploitable fitness trade-offs. However, our study has only characterized a single phage-resistant isolate, each, from AB900 and A9844, and there exists the potential for other independently evolved resistance mutations to emerge that lack these phenotypes. Thus, a key question is whether the evolution of phage-resistance in *A. baumannii* is repeatable and predictable. An analysis of the K loci in heterogeneous populations of phage-resistant mutants could demonstrate the diversity of mutations occurring therein or, conversely, point to other pathways of phage-resistance evolution not involving the K locus. Finally, an important follow-up experiment should aim to identify whether the mechanisms of phage-resistance acquisition reported here also occur *in vivo.*

We previously theorized an approach to phage therapy delivery based on the exploitation of phage-resistance (17). As demonstrated in this study, the emergence of phage-resistance in *A. baumannii* can be accompanied by exploitable trade-offs. But in order to consistently and reliably take advantage of them, we require a detailed characterization of the phages, their hosts, their receptors and their mechanisms of infection and resistance. Ideally, this approach to phage therapy would employ phages that exhibit strong antimicrobial effects, and force a predictable evolutionary response by the pathogen, through known mechanisms of phage-resistance, thereby allowing physicians to foresee their next therapeutic intervention and stay a step ahead of the pathogen (Fig. 6). In a previous clinical case of phage therapy use against MDR *A. baumannii* (22), where the trade-offs of capsule loss and resensitization to an antibiotic were seen in phage-resistant mutants, these findings were not anticipated nor mechanistically explained, arguably limiting their efficient clinical exploitation.

The ultimate goal of our research is to enable the successful clinical translation of phage therapy against *A. baumannii* and other MDR pathogens. Importantly, the findings of our study highlight that phage therapy should not be proposed as a replacement for antibiotic therapy. In fact, we demonstrate that phages can be used not only for their lytic activity, but, when combined with *a posteriori* knowledge of receptors and emergence of phage-resistance, as a resensitization strategy empowering other antimicrobial agents. Improving the efficacy, and furthering the clinical impact of phage therapy, is an important step in the fight against the postantibiotic era.

**Figure 6.**
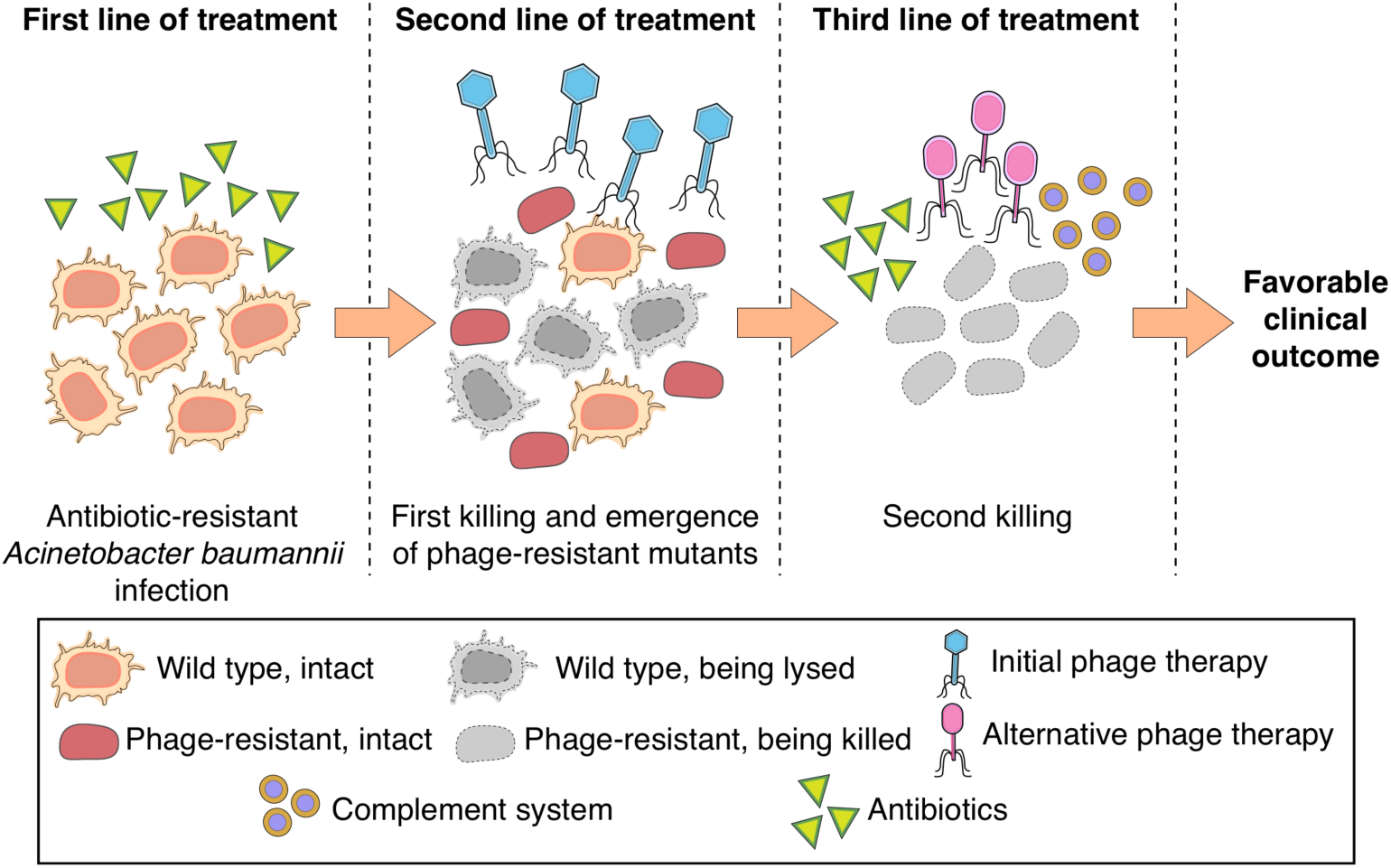
An approach toward phage therapy utilizing lytic activity of phages, *a posteriori* knowledge of receptors, predictable emergence of phage-resistance and exploitation of fitness trade-offs. An MDR *A. baumannii* infection (capsulated red cells) will typically withstand antibiotic treatment (first line of treatment). We can pursue the use of phage therapy (second line of treatment), the lytic activity of the phages resulting in a first killing and the emergence of phage-resistant mutants (unencapsulated red cells). We can exploit the trade-offs of phage-resistance using a third line of treatment that can include re-potentiated antibiotics, alternative phages, and the immune system. This will result in a second killing leading to an improved clinical outcome.

## Materials and Methods

### Bacterial strains, plasmids, and culture conditions

A complete list of strains and plasmids used in this study can be found in Table S2. Bacteria were cultured using either lysogeny broth (LB) or Mueller Hinton broth (MHB) (Sigma-Aldrich, Australia), at 37 °C with aeration, supplemented with agar and/or 50 μg/ml of kanamycin sulfate, as required.

### Phage isolation, amplification and storage

Phages were isolated from raw sewage samples obtained from several states in Australia (Queensland, New South Wales, and Victoria). Aliquots of sewage were combined with overnight cultures of up to three bacterial strains, and supplemented with 10x LB and 10 mM CaCl_2_ and MgSO_4_. Mixtures were incubated overnight and the resulting lysates were purified with the previously described Phage-on-Tap protocol (36). The lysates were titrated in plaque forming units (PFU) per ml and stored at 4 °C.

### Determination of phage host range and efficiency of plating

The lytic activity of each phage was screened against nine *A. baumannii* strains with the standard spot assay as previously described (56). For comparing the efficiency of phage infection between hosts, the efficiency of plating (EOP) assay was used. Overnight bacterial cultures were standardized to an optical density at 600 nm (OD_600_) of 0.3 using a spectrophotometer. This density was previously assessed to be roughly equivalent to 5×10^7^ CFU/ml. Next, soft agar overlays were set up with known concentrations of phage lysate (10^1^-10^3^ PFUs). After incubation, the plates were inspected for plaques and the average number of plaques was recorded and compared as a percentage of the plaques found in the original host of isolation.

### Transmission Electron Microscopy (TEM)

Further purification of the phage lysates, using the Vivaspin 6 centrifugal concentrator (MWCO 1,000,000 kDa) (Sartorius, Australia), was required for high-quality electron microscopy visualization. Then, 10 µl droplets of phage suspension were placed on copper TEM grids (200 mesh, SPI, USA) with carbon-coated ultrathin (invisible on the water surface) formvar film (57). They were left for 30 seconds and then dried using filter paper. A 10 µl droplet of uranyl acetate water solution (1% w/v) was then added to the grid surface and left for 20 seconds. The grid was dried with filter paper, and then examined under a TEM (JEM-1400 Plus; Jeol, Japan) at an accelerating voltage of 80 kV.

### One-step phage killing curves and adsorption assay

Bacteria from overnight cultures and phages from pure lysates were mixed at a multiplicity of infection (MOI) of 0.01 (10^6^ PFU/ml to 10^8^ colony forming units [CFU]/ml) in LB. The suspensions were incubated at 37 °C with aeration. At each timepoint, samples of the suspensions were transferred into chloroform-saturated PBS, vortexed, and then centrifuged at 3,500x g for 3 min. The supernatant was diluted in PBS and plated in duplicate for quantification of free phage particles.

### Extraction of phage DNA, minION sequencing and phage genome analysis

A starting volume of 1.8 ml of phage lysate with a concentration of > 10^10^ PFU/ml was treated with DNase and RNase A. DNA was extracted using the phenol-chloroform method and purified with isopropanol and sodium acetate as described previously (58). The extracted DNA was dissolved in nuclease free water, then quantified and its purity assessed using a Nanodrop (NanoDrop Technologies, Wilmington, DE, USA), Qubit fluorometer (Life Technologies, Carlsbad, CA, USA), and 1% agarose gel electrophoresis. 1 μg of DNA was used for library preparation using the Ligation Sequencing Kit 1D (SQK-LSK109, Oxford Nanopore Technologies, ONT, Oxford UK) following the manufacturer’s instructions. The sequencing libraries were multiplexed and loaded into the Flow Cell FLO-MIN106 Spot-ON of the MinION system using a Library Loading Bead Kit R9 according to the manufacturer’s instructions. The raw reads were obtained using MinKNOW v1.7.14 in a 48 h-run experiment and base calling was performed using ONT default software. Deepbinner v0.2.0 (59) was used to demultiplex the raw reads. Reads smaller than 500 bp were removed using Trimmomatic v.039 (60) and the remainder assembled using Unicycler v0.4.3 (61). Genome annotation was performed by combining multiple tools: the RAST server (62), PHASTER (63) and VIBRANT (37).

### Lysogen isolation

Based on a previously described protocol (64), a spot assay was established with the phage-host pair of interest. After 48 h of incubation, cloudy and confluent centers of bacterial growth within zones of lysis were identified. Bacterial growth from these zones was scraped off using a sterile loop and streak-plated. After overnight incubation, 36 colonies were tested for lysogeny using a patch test. For this test, single colonies were patch-plated onto two LB plates, the second one of which contained a soft agar lawn of the wild type host. After 48 h of incubation, the plates were examined looking for halos of lysis around the patches of the second plate. All presumptive lysogens then underwent three rounds of single-colony purification before repeating a patch test. Finally, liquid cultures of the possible lysogens were pelleted, and the supernatants were filtered through a 0.2 μm filter and tested for spontaneous phage release using spot assays.

### Bacterial kinetics

Growth kinetics were determined using a 96 well plate format, measuring OD_600_ every 10 min with a starting inoculum of 10^6^ CFUs, unless otherwise specified, and phage PFUs at MOIs of either 0, 1 or 0.1. Where required, area under the curve (AUC) was calculated from a baseline of y = 0.1 and the percent difference calculated (65).

### Isolation of phage-resistant *A. baumannii* mutants

Following the same principle as the spot assay, 25 μl aliquots of pure lysate (concentration > 10^8^ PFU/ml) were spotted onto bacterial lawns. After an overnight incubation, bacterial colonies growing in the middle of lysis zones were picked and streaked onto LB agar plates for two rounds of single-colony isolation. The phage-resistant phenotypes were confirmed using three methods: inverted spot plate assays (66), standard soft agar overlay assays, and *in vitro* growth curves. Phage-resistant mutant strains were tested bimonthly to ensure reversion to a phage-sensitive phenotype did not occur.

### Extraction of bacterial genomic DNA, sequencing, bacterial genome comparison and phylogenetic analysis

The GenElute™ Bacterial Genomic DNA Kit protocol (Sigma-Aldrich, Australia) was used for DNA extraction. Bacterial genomic DNA was tested for purity as previously described for phage DNA and vacuum dried into a pellet for transport. Sequencing was performed using the Illumina® HiSeq 150 bp paired-end platform at the Genewiz® facilities (Suzhou, China).

The genomes of AB900, A9844 and their phage-resistant counterparts, were independently assembled from the paired-end 150 nt Illumina reads. Adapter sequences were clipped using Trimmomatic v0.39 (60) in palindrome mode, with quality filtering, quality trimming and length filtering (AVGQUAL:35 SLIDINGWINDOW:5:28 MINLEN:80). Trimming outcomes were checked in trimviz (github.com/MonashBioinformaticsPlatform/trimviz). Reads were assembled using Spades v3.12.0 (67) with a coverage cutoff of 10 in ‘careful’ mode. To identify variants, the filtered sequencing data was mapped back onto the assemblies using BWA (68) as implemented in RNAsik v1.5.0 (69). Both phage-sensitive (S) and phage-resistant (R) sequencing data was mapped back onto each of the genome assemblies from S and R isolates, resulting in 4 alignment files per strain pair (S on S; S on R; R on S; and R on R). Pileup files were generated using samtools mpileup v1.8 and processed using Varscan2 v2.4.0 (70) in both “pileup2snp” and “pileup2indel” modes. Candidate variants (several thousand per alignment) were then filtered using a custom Python script for: 1) contig length > 500 nt; 2) contig coverage in the Spades assembly > 50; 3) realignment coverage > 30 reads at the variant site; 4) if the variant is also detected in the self-alignment (S on S) its frequency (proportion of supporting reads vs total overlapping reads) must be at least 25% higher in the non-self-alignment (R on S). Surviving variants were few enough in number to be manually verified. R on S alignments around candidate variants were viewed in IGV v2.3.59 (71) together with the .gff3 annotation files generated by Prokka v.1.13.7 (72) to check for coding-region disruptions, and compared to the self-alignment (S on S) to check for alignment artefacts. The reciprocal alignments (S on R, checking R on R for alignment artefacts) were also verified in the homologous region of the R assembly.

To infer overall sequence-based relationships between the strains in this study, the publicly available genome assemblies were downloaded (Table S3). For strain AYP A22, however, only raw sequencing data was available, so this was trimmed and assembled as above (Trimmomatic parameters were adjusted to AVGQUAL:28 SLIDINGWINDOW:5:23 MINLEN:40 without adapter-trimming). All assemblies were *de novo* annotated using Prokka and core-genome alignments of coding regions common to all the strains were created using Roary v3.11.2 (73). Alignments were converted to PHYLIP format and fed into RAXML v8.2.12 (74) (with the GTRCAT substitution model). To illustrate the relationships between the strains documented in Figure 1, the data was visualized using the R package APE v5.3 (75).

### Gene knockout and complementation

All PCR reactions mentioned in this section were performed using KAPA HiFi DNA Polymerase (Hot Start and Ready Mix formulation) (Roche, Switzerland). Genes of interest were disrupted using a variation of the methods described previously (76). Briefly, ∼1000 bp upstream and downstream of the target genes were PCR-amplified from genomic DNA. The kanamycin resistance cassette was amplified from pKD4 (76), and three products combined by splice overlap PCR using primers specified in Table S4. The linear constructs were gel purified using a PCR cleanup kit (Sigma-Aldrich, Australia), confirmed by Sanger sequencing and concentrated to ∼500 ng/μl.

*A. baumannii* cells were made electrocompetent as follows. Overnight cultures were diluted 1:20 and allowed to grow to an OD_600_ of 0.7. A volume of 5 ml of culture per transformation was pelleted by centrifugation, the cells were washed 3 times in 10% (v/v) glycerol and finally resuspended in 10% glycerol. The cell suspension was mixed with recombinant DNA (∼3 μg) and incubated at room temperature for 15 min. The mixture was transferred to a sterile 2 mm electroporation cuvette (Biorad, Australia) and immediately pulsed with a Bio-Rad Micropulser (EC3, 3.0 kV). Pre-warmed (37 °C) Super Optimal Broth (SOB) was immediately added and the cells were transferred to a new tube. The cells were recovered for 3 hours rotating at 37 °C. The recovered cells were pelleted, resuspended in SOB, and spread-plated onto LB agar with and without 50 μg/ml of kanamycin. Isolated colonies from the transformation were patch-plated on LB agar containing 50 μg/ml of kanamycin for two successive re-streaks to ensure kanamycin resistance. Colonies of the knockout mutants were PCR-amplified with the primers on Table S4, ran on a 0.7% agarose gel, and the bands were gel-purified and Sanger-sequenced in order to identify the successful disruption of genes of interest via the insertion of the kanamycin resistance cassette.

For complementing the genes of interest back into the knockout mutants, the genes were PCR-amplified using primers containing cut sites for the restriction enzymes *EcoR*I (forward) and *Sal*I (reverse) (New England Biolabs, NEB, Australia) for *gtr29*, or *Sal*I exclusively in the case of *gpi*. Next, two overnight double-digestions were set up at 37 °C for the plasmid pBAD18Kan-Ori (77) and the genes of interest using the restriction enzymes. The digested products were run on a 0.7% agarose gel, excised, purified and ligated for 2 h at room temperature using the Instant Sticky-end Ligase Master Mix (NEB, Australia) following the manufacturer’s protocol. The ligated product was then chemically transformed into *Escherichia coli* DH5-*α* competent cells (NEB, Australia) following the company’s protocol. The transformed colonies were patch-plated onto kanamycin-LB plates (50 μg/ml), and confirmed via Sanger sequencing. These confirmed colonies underwent plasmid prep using the GenElute™ Plasmid Miniprep Kit (Sigma-Aldrich, Australia) and the complement-containing plasmid was electroporated into the knockout mutant as per the protocol above.

### Capsule quantification and observation

Quantification of surface polysaccharides was performed following the published protocol of the sulfuric acid-phenol reaction (39). For qualitative observation of capsule thickness, a Maneval’s capsule stain was performed using a bacterial colony mixed with one drop of Congo Red (1% aqueous) on a clean glass slide. After drying, the slide was flooded with Maneval’s modified stain for 1 min, washed with water and blotted dry. Slides were observed using light microscopy under oil immersion. For scanning electron microscopy (SEM) imaging of the surface of bacterial hosts, a 20 μl droplet of 5 times-PBS washed bacterial suspension was placed onto a fresh gold-coated silicon wafer (5 ϗ 5 mm) and the cells allowed to settle for 5 min, followed by washing with PBS. The wafer was then placed in 2.5% Glutaraldehyde in PBS solution for 15 min, washed with deionized water and dehydrated by immersing in increasing concentrations of ethanol for 3 minutes each. Residual ethanol was removed with a critical point dryer CPD 030 (BAL-TEC AG, Liechtenstein). The sample was mounted on a standard metal SEM stub and then coated with a ∼10 nm thick gold layer using a sputter coater SCD 005 (BAL-TEC AG, Liechtenstein). The samples were examined under high vacuum within the FE-SEM ThermoFisher Elstar G4 at an accelerating voltage of 2 kV, secondary electron mode (SE), and a working distance of 4 mm, operating in immersion mode with the through lens detector (TLD).

### Biofilm formation assay

48 h biofilm formation on polystyrene 96-well plates was assessed using crystal violet staining and spectrophotometry as previously described (78).

### Serum killing assay

From overnight cultures of the strains, a 1:100 subculture was made and incubated to reach mid-exponential phase (approximately 2-3 h). Meanwhile, the working volume of human serum (Sigma-Aldrich, Australia) was divided in half, storing one half at 4 °C and heat-inactivating the other half at 56 °C for 30 min. After incubation, the cells of each culture were harvested via centrifugation, washed, and resuspended in PBS, then standardized to an OD_600_ of 0.3. Active or heat-inactivated human serum was then diluted in PBS to a 50% solution. In a glass test tube, 50% serum/PBS was inoculated with 10^5^ CFU of bacteria, mixed and incubated. At regular intervals, samples from each culture were extracted to perform serial dilutions and CFU counts. *E. coli* DH5-*α* was used as a control of serum activity, as it is highly susceptible to human serum but resistant to heat-inactivated serum.

### Antibiotic susceptibility assay

Minimum inhibitory concentrations (MICs) of 7 antibiotics were assessed using the microbroth dilution protocol, as previously described (79) and interpreted using the cut-off values from the Clinical and Laboratory Standard Institute. The antibiotics used were: ampicillin/sulbactam (SAM), ceftazidime (CFZ), imipenem (IPM), gentamicin (GEN), amikacin (AMK), minocycline (MIN), and ciprofloxacin (CIP) (Sigma-Aldrich, Australia). Size of the bacterial inoculum for each test was standardized at 10^5^ CFU, in a volume of 200 μl per test. *E. coli* strains ATCC 25922 and ATCC 35218 were used as quality controls for each batch of microbroth dilution tests.

### Animal experiments and animal ethics

Four female, 6-to-10 weeks old, BALB/c mice per group were used. Bacterial inoculums of either wild type or phage-resistant *A. baumannii* were prepared to a concentration of 10^6^ CFU in 100 μl of PBS. Before administration, the bacterial suspensions were mixed in a 1:1 ratio with 6% porcine stomach mucin (Sigma-Aldrich, Australia) in PBS, for a total volume of 200 μl. Mice were injected intraperitoneally and followed for up to 8 h. At the humane endpoint, blood was extracted through cardiac puncture and a laparotomy was performed to obtain a section of the liver, the right kidney, and spleen. The organs were weighed and then homogenized in PBS. Blood and organ suspensions were then serially diluted in PBS (from 10^−1^ to 10^−5^) and the bacterial burden quantified by CFU counting, and normalized by organ weight. All protocols involving animals were reviewed and approved by the Monash University Animal Ethics Committee (Project ID: E/1689/2016/M) and complied with the National Health and Medical Research Council guidelines. Animals were housed at the Monash Animal Research Facility, Monash University.

### Graphing and statistics

Graphing and statistical analyses were performed with GraphPad Prism 7 (GraphPad Software, Inc.). All *in vitro* experiments were performed in triplicate, with at least two technical replicates each. Where appropriate, scatter plots and medians were presented (80). Both parametric and nonparametric statistical analyses were performed, and the threshold value of two-tailed p < 0.05 used for statistical significance.

## Supporting information

FG02 Annotation

CO01 Annotation

## Acknowledgements

We acknowledge expert advice and input from Laura Perlaza-Jiménez and Rhys Dunstan in the early stage of this project. We also gratefully acknowledge the use of facilities within the Monash Ramaciotti Cryo EM platform, as well as Dr Alex Fulcher from Monash Micro Imaging, and A/Prof Alex de Marco for granting access to the SEM. Christian Vásconez provided feedback regarding the readability, clarity, and structure of the manuscript.

Fernando L. Gordillo Altamirano acknowledges the support received from Monash University through the Monash Postgraduate Research Scholarships funding his doctoral studies. Trevor J. Lithgow is an ARC Australian Laureate Fellow (FL130100038). This work, including the efforts of Jeremy J. Barr, was funded by the National Health and Medical Research Council (NHMRC: 1156588), and the Perpetual Trustees Australia award (2018HIG00007).

## Author Contributions

Conceptualization: FGA, TJL, AYP, JJB; Methodology: FGA, JHF, RP, XK, MT, SA, FM, DK, JJB; Formal Analysis: SA; Investigation: FGA, JHF, RP, XK, MT, DS, CO, LK, DK; Resources: MKO’B, TJL, AYP, JJB; Writing – Original Draft Preparation: FGA, JJB; Writing – Review and Editing: FGA, JHF, RP, XK, MT, DS, SA, FM, CO, LK, DK, MKO’B, TJL, AYP, JJB; Supervision and Funding Acquisition: TJL, AYP, JJB.

## Supplementary Information

**Figure S1.**
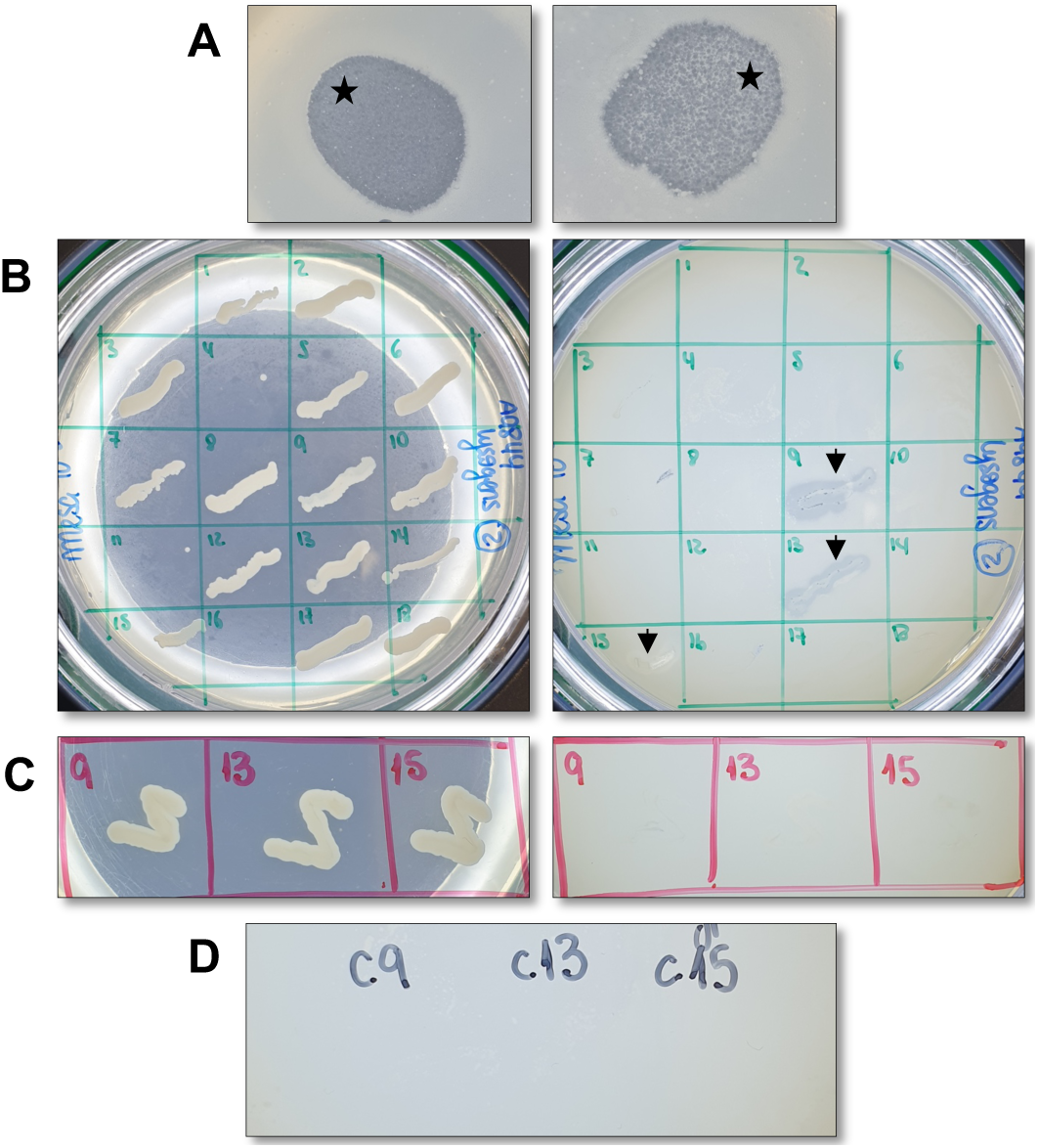
Attempted production of A9844 lysogens. A: Spot assay with droplets containing 10^6^ (left) and 10^4^ (right) PFU of øCO01 after 48 h of incubation. Black stars denote the zones of lysis containing cloudy bacterial growth. B: Patch assay. Bacterial growth scraped off from the lysis zones was streaked-plated and individual colonies were patched onto LB (left) and LB with an A9844 overlay (right) plates. Black arrows (colonies 9, 13 and 15) show bacterial growth with surrounding lysis, indicating the presence of phage øCO01, possibly released from a lysogen. Panel only shows 18/36 screened colonies. C: After three rounds of single-colony purification, the patch test was repeated for colonies 9, 13 and 15, without indication of phage presence. D: Spot assay performed with the filtered supernatant from cultures of colonies 9, 13 and 15, with no indication of spontaneous phage release.

**Figure S2.**
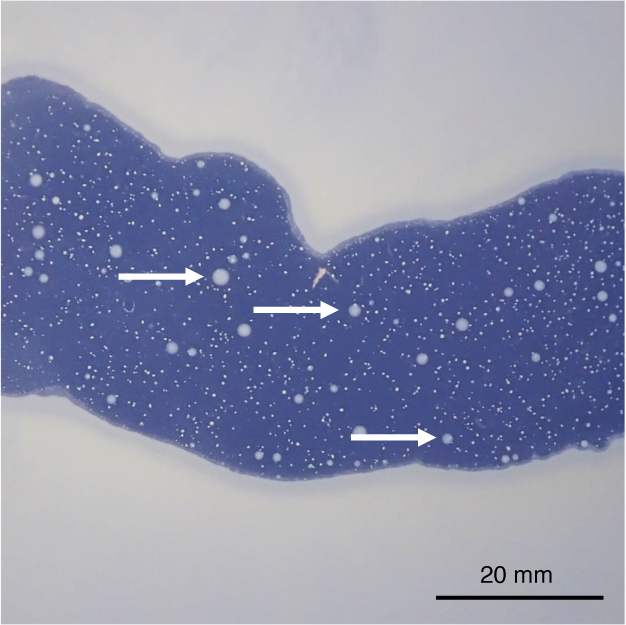
Spot assay of strain A9844 with phage øCO01, showing the large central lysis zone, with several bacterial colonies growing within (white arrowheads). These colonies were assumed to be phage-resistant and underwent the pipeline for confirming this phenotype.

**Figure S3.**
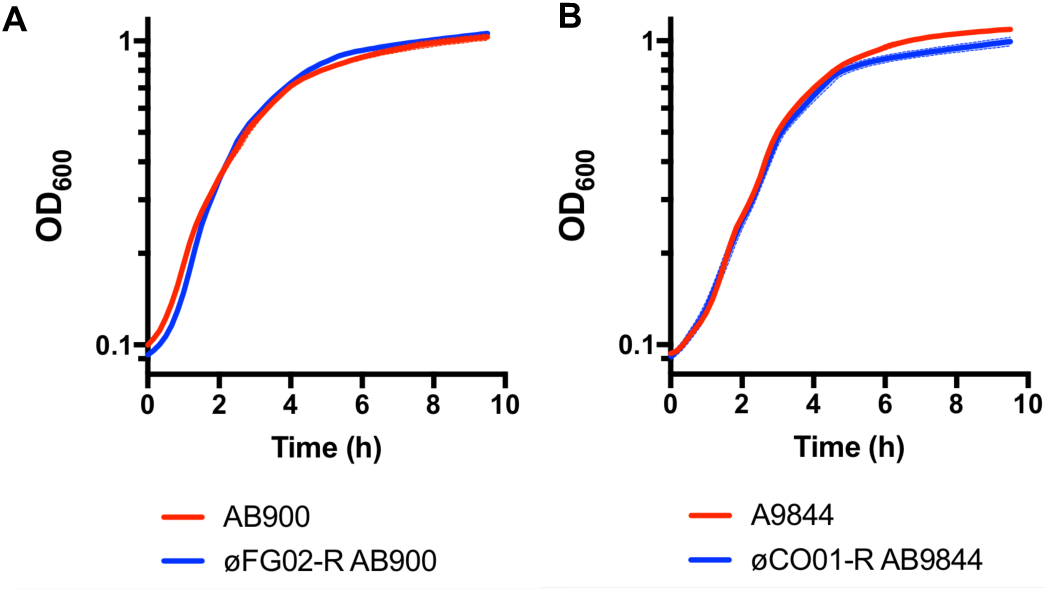
Growth curves of A. baumannii strains AB900 and A9844 (red) and their phage-resistant mutants øFG02-R AB900 and øCO01-R A9844 (blue) in the absence of phages. Shaded zones represent SD (n = 3).

**Figure S4.**
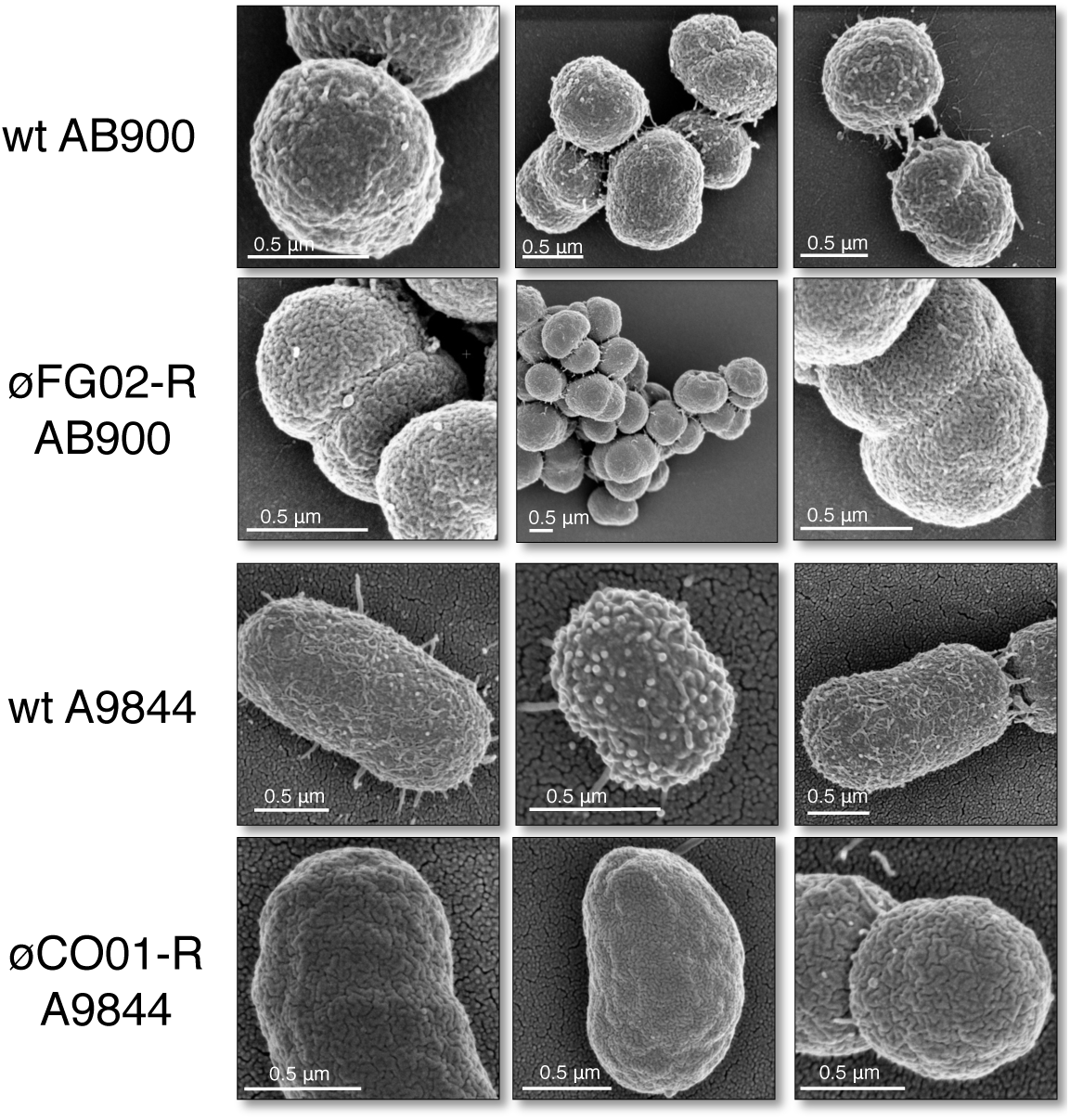
Additional scanning electron microscopy images of the wild type and phage-resistant strains in this study.

**Figure S5.**
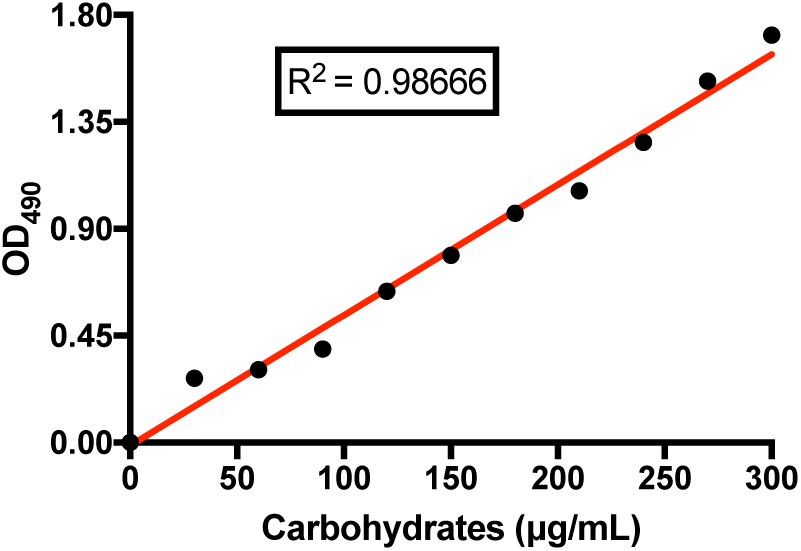
Standard curve for the capsule quantification assay. Carbohydrate standards were prepared by diluting a carbohydrate stock solution (50:50 mixture of 0.5 mg/ml each of sucrose and fructose) into 1 ml aliquots ranging from 0 to 300 μg/ml of carbohydrate using distilled water. The standards were treated with the phenol-chloroform reaction, and the absorbance measured at OD_490_.

**Table S1.** Raw data of minimum inhibitory concentrations (MIC) in μg/ml from the antibiotic resistance panel of AB900, A9844, øFG02-R AB900 and øCO01-R A9844. MICs determined via the microbroth dilution method.

|  | AB900 |  |  | øFG02-R AB900 |  |  |
| --- | --- | --- | --- | --- | --- | --- |
|  | Replicates |  |  | Replicates |  |  |
|  | 1 | 2 | 3 | 1 | 2 | 3 |
| Ampicillin + Sulbactam | 2/1 | 8/4 | 4/2 | 2/1 | 4/2 | 2/1 |
| Ceftazidime | >32 | >32 | 8 | 2 | 2 | 1 |
| Imipenem | 0.12 | 0.12 | 0.25 | ≤0.06 | ≤0.06 | 0.25 |
| Ciprofloxacin | 0.5 | 0.25 | 0.25 | 0.25 | 0.12 | 0.25 |
| Amikacin | 2 | 1 | 4 | ≤0.25 | 1 | 4 |
| Gentamicin | 0.5 | 0.5 | 1 | 0.5 | 0.5 | 0.5 |
| Minocycline | ≤0.12 | ≤0.12 | 0.25 | ≤0.12 | ≤0.12 | 0.25 |
|  | A9844 |  |  | øCO01-R A9844 |  |  |
|  | Replicates |  |  | Replicates |  |  |
|  | 1 | 2 | 3 | 1 | 2 | 3 |
| Ampicillin + Sulbactam | 16/8 | 16/8 | 16/8 | 8/4 | 8/4 | 16/8 |
| Ceftazidime | >16 | 64 | 64 | >16 | 32 | 64 |
| Imipenem | 8 | >32 | 32 | 8 | 16 | 32 |
| Ciprofloxacin | >8 | >8 | >8 | >8 | >8 | >8 |
| Amikacin | 0.25 | ≤0.25 | 2 | 0.5 | 2 | 0.25 |
| Gentamicin | 2 | 1 | 0.5 | 2 | 0.5 | 2 |
| Minocycline | 2 | 2 | 2 | 2 | 1 | 1 |

**Table S2.**
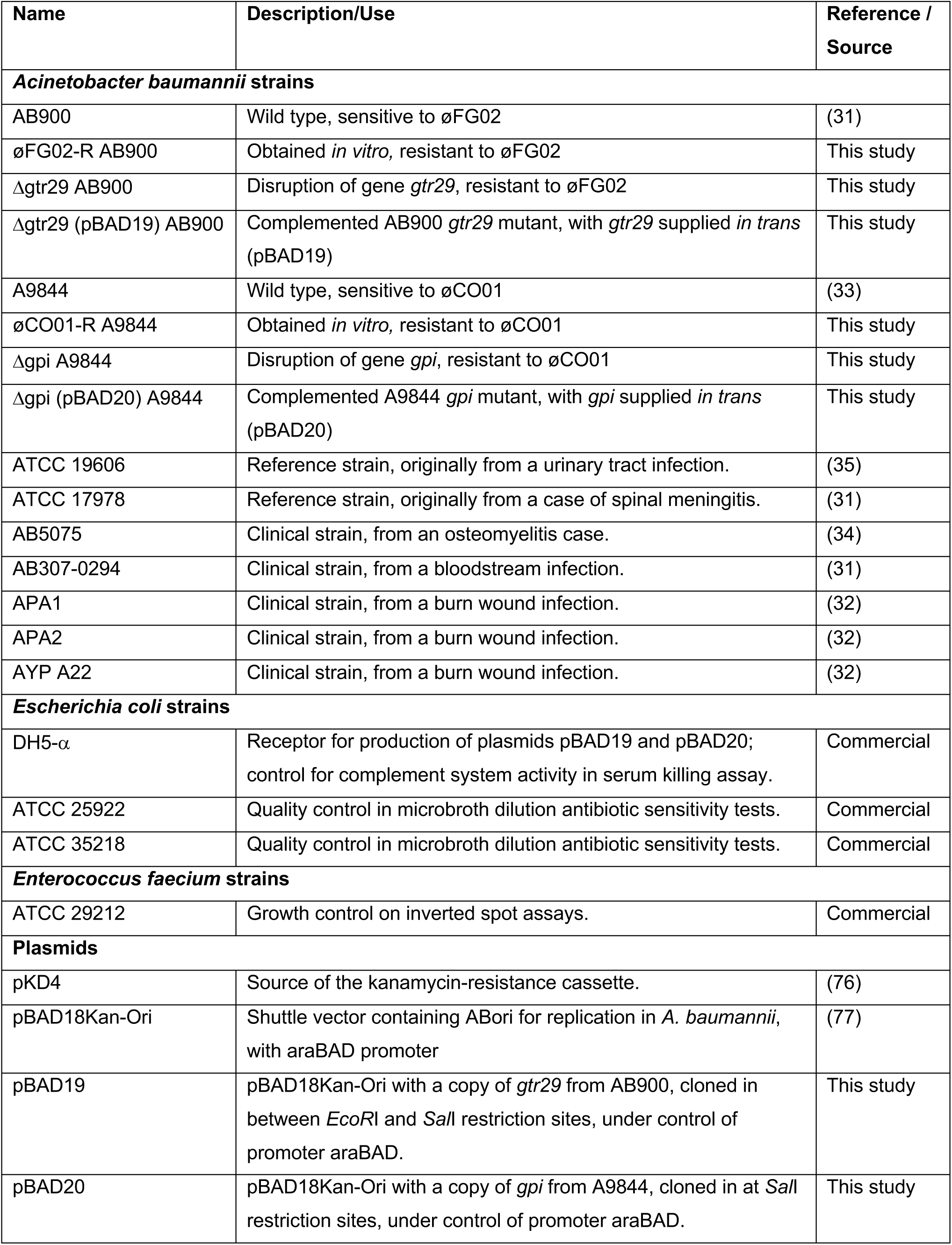
Strains and plasmids used in this study

| Name | Description/Use | Reference / Source |
| --- | --- | --- |
| ***Acinetobacter baumannii* strains** |  |  |
| AB900 | Wild type, sensitive to øFG02 | (31) |
| øFG02-R AB900 | Obtained *in vitro*, resistant to øFG02 | This study |
| Δgtr29 AB900 | Disruption of gene *gtr29*, resistant to øFG02 | This study |
| Δgtr29 (pBAD19) AB900 | Complemented AB900 *gtr29* mutant, with *gtr29* supplied *in trans* (pBAD19) | This study |
| A9844 | Wild type, sensitive to øCO01 | (33) |
| øCO01-R A9844 | Obtained *in vitro*, resistant to øCO01 | This study |
| Δgpi A9844 | Disruption of gene *gpi*, resistant to øCO01 | This study |
| Δgpi (pBAD20) A9844 | Complemented A9844 *gpi* mutant, with *gpi* supplied *in trans* (pBAD20) | This study |
| ATCC 19606 | Reference strain, originally from a urinary tract infection. | (35) |
| ATCC 17978 | Reference strain, originally from a case of spinal meningitis. | (31) |
| AB5075 | Clinical strain, from an osteomyelitis case. | (34) |
| AB307-0294 | Clinical strain, from a bloodstream infection. | (31) |
| APA1 | Clinical strain, from a burn wound infection. | (32) |
| APA2 | Clinical strain, from a burn wound infection. | (32) |
| AYP A22 | Clinical strain, from a burn wound infection. | (32) |
| ***Escherichia coli* strains** |  |  |
| DH5-α | Receptor for production of plasmids pBAD19 and pBAD20; control for complement system activity in serum killing assay. | Commercial |
| ATCC 25922 | Quality control in microbroth dilution antibiotic sensitivity tests. | Commercial |
| ATCC 35218 | Quality control in microbroth dilution antibiotic sensitivity tests. | Commercial |
| ***Enterococcus faecium* strains** |  |  |
| ATCC 29212 | Growth control on inverted spot assays. | Commercial |
| **Plasmids** |  |  |
| pKD4 | Source of the kanamycin-resistance cassette. | (76) |
| pBAD18Kan-Ori | Shuttle vector containing A<sub>Bori</sub> for replication in *A. baumannii*, with araBAD promoter | (77) |
| pBAD19 | pBAD18Kan-Ori with a copy of *gtr29* from AB900, cloned in between *EcoRI* and *SaII* restriction sites, under control of promoter araBAD. | This study |
| pBAD20 | pBAD18Kan-Ori with a copy of *gpi* from A9844, cloned in at *SaII* restriction sites, under control of promoter araBAD. | This study |

**Table S3.**
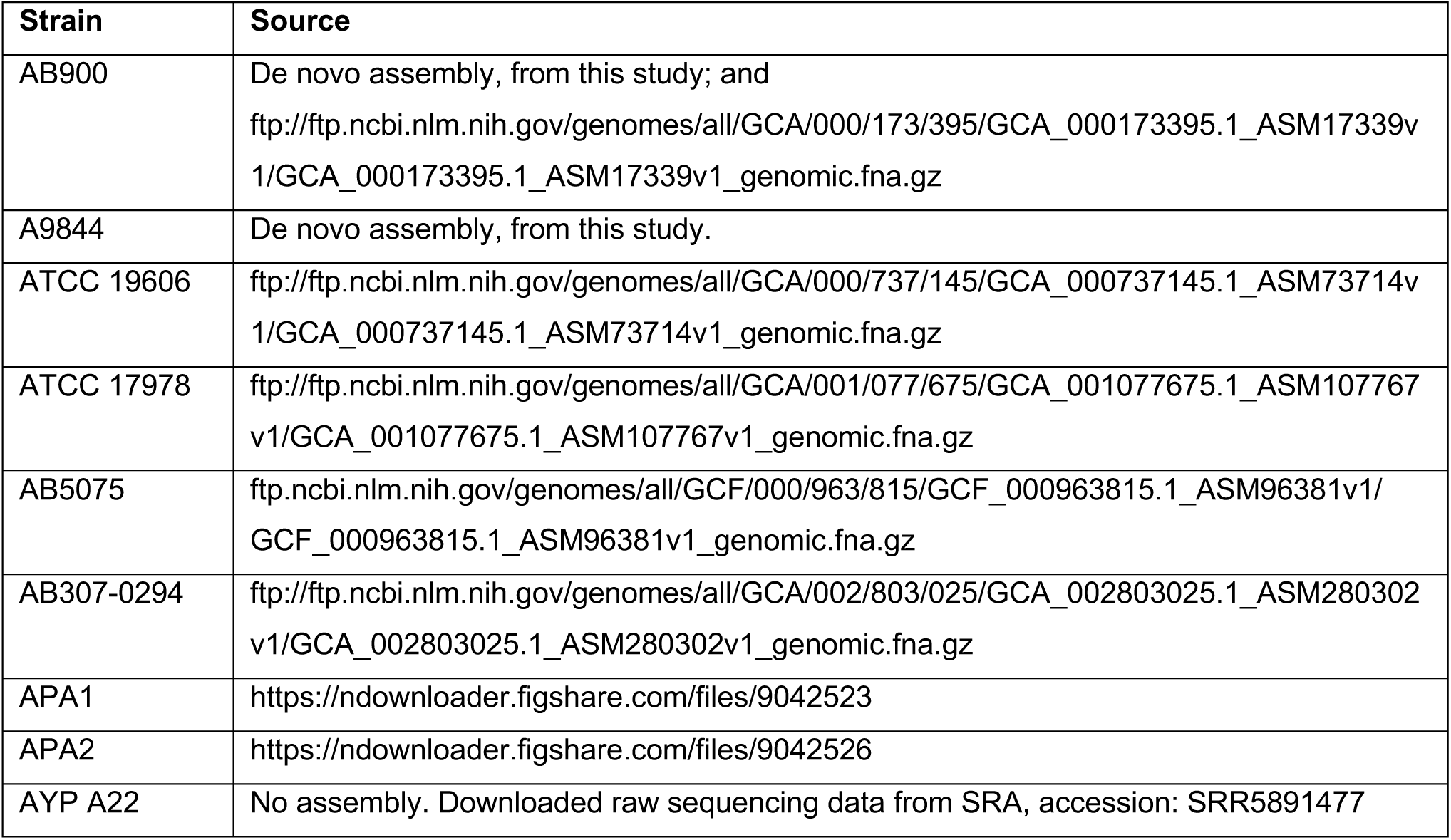
Genome assembly sources for the phylogenetic analysis of *A. baumannii* strains.

**Table S4.**
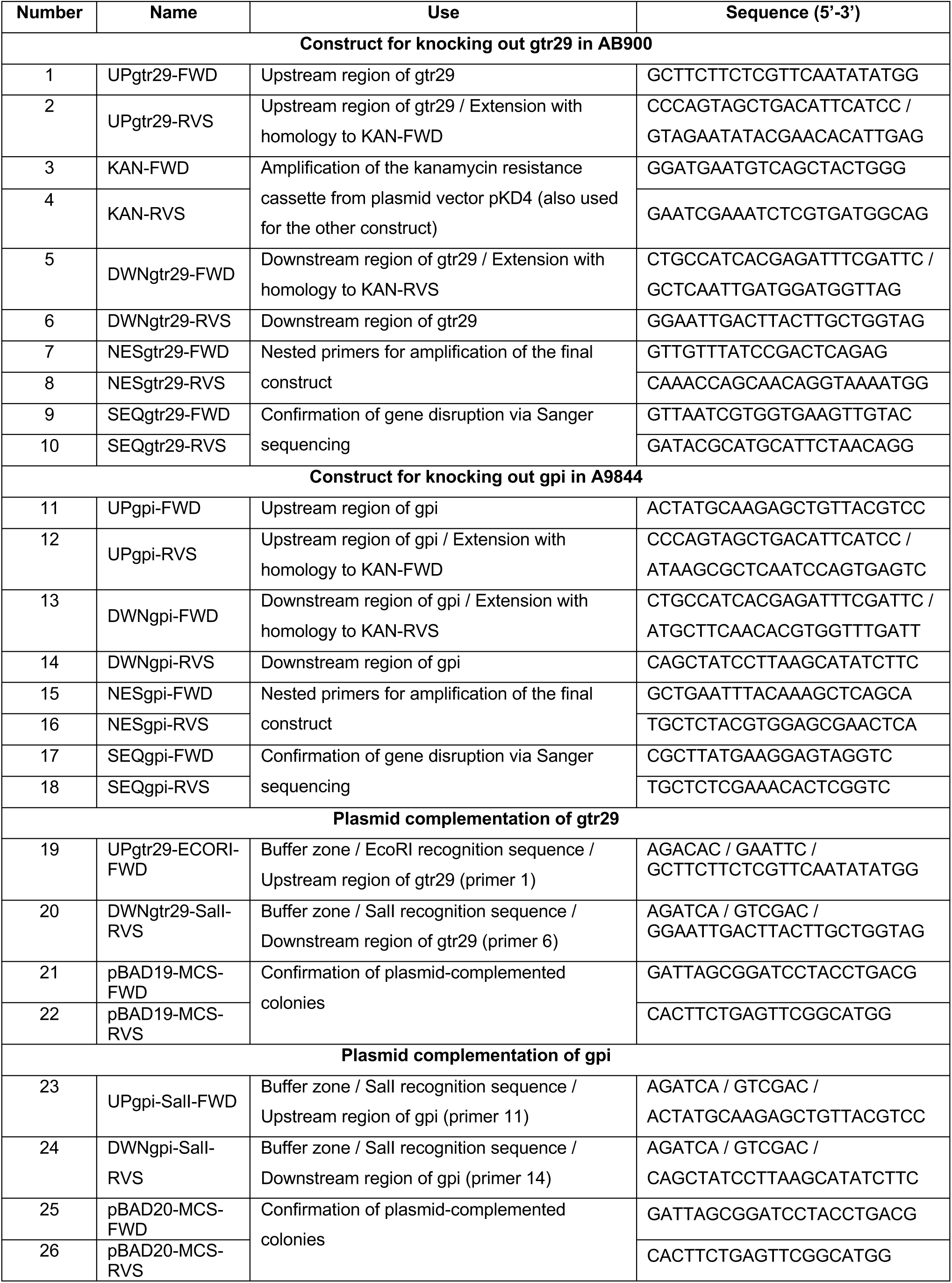
Primers used in the genetic engineering of *A. baumannii* strains AB900 and A9844 in this study. ‘/’ indicates the addition of an extension sequence.

